# Integrated genomic and proteomic analysis of the mouse-adapted *Staphylococcus aureus* strain JSNZ

**DOI:** 10.1101/2025.09.03.674026

**Authors:** Hannes Wolfgramm, Larissa Milena Busch, Jöran Tebben, Henry Mehlan, Lisa Hagenau, Thomas Sura, Tilly Hoffmüller, Elisa Bludau, Manuela Gesell Salazar, Alexander Reder, Stephan Michalik, Leif Steil, Kristin Surmann, Ulrike Mäder, Silva Holtfreter, Uwe Völker

## Abstract

Mouse-adapted *Staphylococcus aureus* strains have become increasingly relevant in infection research thanks to their ability to better recapitulate clinical infection dynamics in mouse models. However, detailed characterisations required to establish a corresponding reference strain are still lacking. The mouse-adapted CC88 strain JSNZ appears to be an ideal candidate for a reference strain, because CC88 is widespread among laboratory mice and frequently employed in mouse colonisation and infection models. Moreover, JSNZ demonstrates high genetic transformability comparable to that of commonly used laboratory strains. Here, we present a comprehensive genomic and proteomic characterisation of JSNZ. Whole genome sequencing was performed using a combination of short and long reads. Proteomic profiling was conducted under standard laboratory conditions in TSB and RPMI during exponential and stationary growth using LC-MS/MS. The updated, closed genome sequence of JSNZ was integrated into *Aureo*Wiki for user-friendly access and direct comparison to long-established reference strains. Genome data revealed a deletion in the restriction endonuclease gene *hsdR*, likely explaining the observed efficient transformation while retaining DNA modification capabilities. This positions JSNZ as a hub for genetic modification of other CC88 isolates. Proteomic profiling of JSNZ indicated broad similarity to common *S. aureus* reference strains. However, a striking exception was the novel serine protease Jep, which constituted approximately 75% of the exoproteome in stationary TSB cultures. Overall, these findings affirm JSNZ as a robust and genetically tractable model strain for murine *S. aureus* infection research and contribute a valuable standardised resource to enhance experimental reproducibility and cross-study consistency in the field.

**Graphical abstract:** 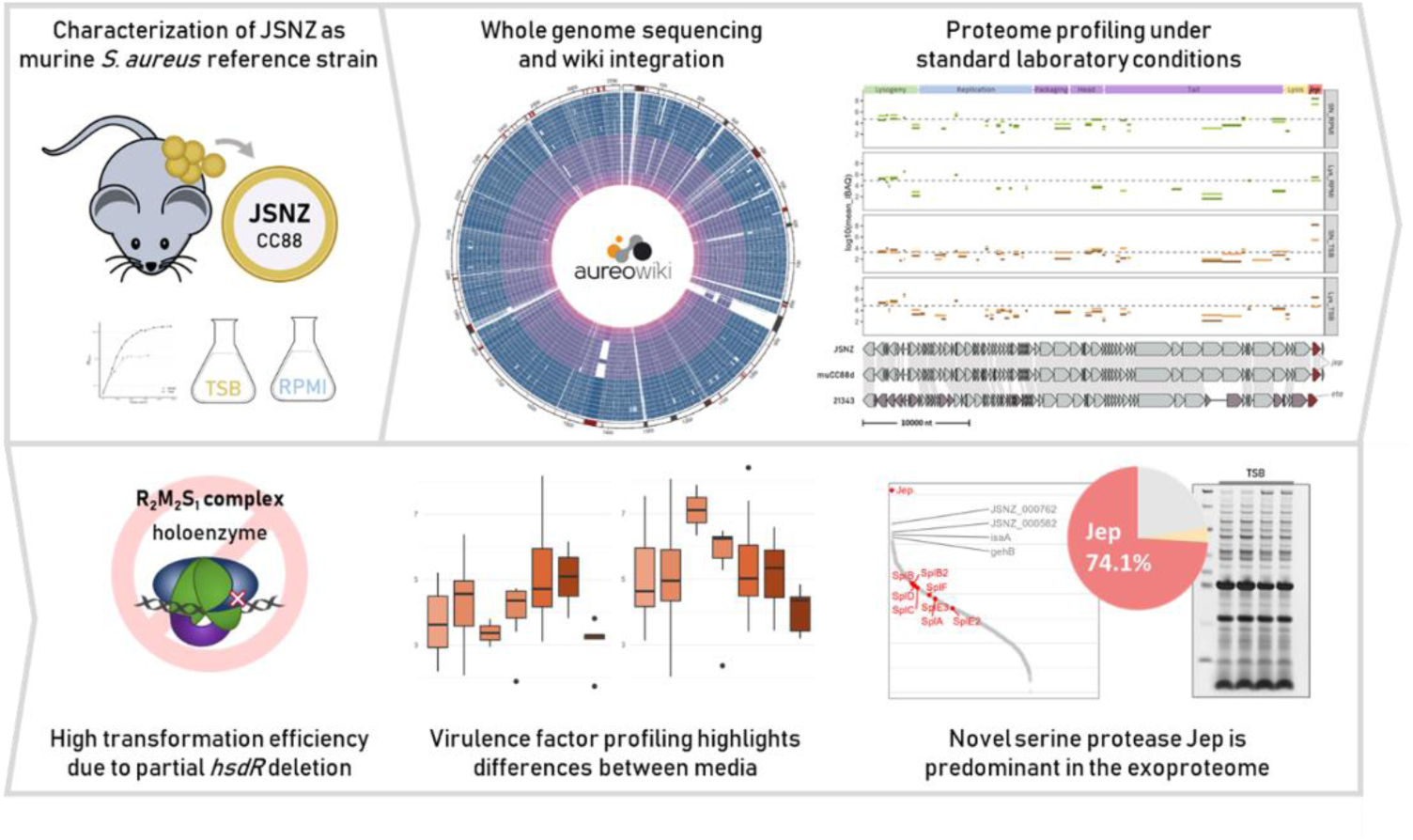

## 1. Introduction

In the race against the continuous adaptation of pathogens, reliable infection models are essential for infection research. The more accurate the model, the more likely it is that the results can be translated into successful therapies (Bulitta *et al*., 2019; Johansen *et al*., 2020; McGonigle and Ruggeri, 2014). Unfortunately, promising therapeutic approaches, such as vaccine candidates, often fail in clinical trials because the employed animal models do not adequately recapitulate the clinical situation (Carey *et al*., 2016; Mrochen *et al*., 2020; Seok *et al*., 2013). This gap is particularly critical for *Staphylococcus aureus*, one of the most relevant human pathogens (Fowler Jr and Proctor, 2014; Ikuta *et al*., 2022; O’Gara, 2017; Salgado-Pabón and Schlievert, 2014). As mice are the primary animal model (not only) for *S. aureus* infection, much of the current knowledge about its virulence and pathogenicity originates from murine infection studies (Kim *et al*., 2014). However, apart from differences in the immune systems of mice and humans (Bjornson-Hooper *et al*., 2022; Mestas and Hughes, 2004; Nauseef, 2023), the commonly used combination of murine host and clinical *S. aureus* isolates imposes several limitations on the model and results in discrepancy compared to human infections: (i) a significantly increased infection dose required to achieve comparable disease phenotypes, (ii) the need for pre-treatment to facilitate initial colonisation, and (iii) a lack of persistent colonisation due to rapid eradication of clinical isolates by the murine host (Holtfreter *et al*., 2013; Kiser *et al*., 1999; Mrochen *et al*., 2020). Each of these steps causes the model to deviate even further from the physiological situation.

The reason for these difficulties arises from the high variability of *S. aureus* strains, which in part exhibit pronounced lineage-specific host tropism and strongly host-adapted virulence factors (Howden *et al*., 2023; Matuszewska *et al*., 2020; Richardson *et al*., 2018). *S. aureus* has a broad host spectrum and colonises many vertebrates, ranging from domestic animals, such as livestock, to wild animals, such as wild rodents and birds (Matuszewska *et al*., 2020). This wide spread is driven by recurrent host switches, accompanied by lineage diversification with sometimes drastic genetic changes to adapt to the new host. In particular, mobile genetic elements (MGE) and the associated virulence factors, ranging from toxins to adhesins and immune evasion factors, as well as resistance patterns play a central role (Haag *et al*., 2019; McCarthy and Lindsay, 2013; Richardson *et al*., 2018; Yebra *et al*., 2022, 2021). Well-documented examples are the pore-forming leukocidins (Boguslawski *et al*., 2020; Koop *et al*., 2017) or the pro-coagulatory proteins coagulase and von Willebrand factor-binding protein (Viana *et al*., 2010; Yebra *et al*., 2024). In consequence, numerous virulence factors of clinical *S. aureus* isolates display reduced activity or are non-functional in mice (Boguslawski *et al*., 2020; Mrochen *et al*., 2020; Salgado-Pabón and Schlievert, 2014).

Mouse-adapted strains are a promising option to circumvent these common issues and improve murine infection models (Boff *et al*., 2025; Trübe *et al*., 2019). Natural *S. aureus* colonisation is found in laboratory mice (Mrochen *et al*., 2018a; Schulz *et al*., 2017) as well as in wild mice and other rodents (Fountain *et al*., 2021; Mrochen *et al*., 2018b; Raafat *et al*., 2020). In recent years, selected mouse-derived isolates have been described and studied in murine infection models. Two of them belong to the most frequent clonal complex (CC) found in laboratory mice, CC88 (JSNZ (Holtfreter *et al*., 2013) and WU1 (Sun *et al*., 2018)), one to the also globally distributed but less frequent CC15 (SaF_1 (Flaxman *et al*., 2017)) and one, isolated from a wild bank vole, belongs to CC49 (DIP (Trübe *et al*., 2019)). Two other isolates were already recovered in the 1990s from spontaneous disease outbreaks in laboratory mice (LS-1 (Bremell *et al*., 1990) and DAK (Kiser *et al*., 1999)), but although they are still in use (Boldock *et al*., 2018; Mohammad *et al*., 2022), no genomic characterisation is available. Of note, the isolates from laboratory mice generally tend to exhibit a colonising character with persistent colonisation, whereas DIP is highly virulent in several murine infection models (Mrochen *et al*., 2020; Trübe *et al*., 2019). Meanwhile, all of these murine isolates are increasingly used in infection studies (e.g. Bobrovskyy *et al*., 2023; Boff *et al*., 2025; Cullum *et al*., 2024; Flaxman *et al*., 2019; Hollwedel *et al*., 2024), with JSNZ currently being the most frequently employed (Chan *et al*., 2024; Clow *et al*., 2020; Fernandes de Oliveira *et al*., 2021; Langley *et al*., 2017; Piewngam *et al*., 2018; Reifenrath *et al*., 2020; Sun *et al*., 2018). Accordingly, strain JSNZ – isolated in New Zealand from preputial gland abscesses of C57BL/6 mice (Holtfreter *et al*., 2013) – might establish itself as a reference strain for optimised murine infection models with mouse-adapted *S. aureus*.

Reference strains are preferably characterised by detailed knowledge of their genome and physiology to support evaluation and interpretation of further studies. In this study, we present an updated and now closed genome sequence for JSNZ. The annotated genome has been enriched with computational regulon predictions and is made available for detailed exploration in *Aureo*Wiki (Fuchs *et al*., 2018), https://aureowiki.med.uni-greifswald.de). Moreover, we examined the proteome and exoproteome profile during exponential and stationary growth in two standard laboratory media used with *S. aureus*: the complex tryptone soy broth (TSB) and the chemically defined Roswell Park Memorial Institute 1640 (RPMI-1640) eukaryotic cell culture medium. The observed proteomic profiles demonstrate the comparability of JSNZ to established reference strains in terms of metabolism and virulence factor profiles, while also suggesting potential features of host adaptation. Notably, the novel secreted serine protease Jep was detected at exceptionally high abundance across all tested conditions, markedly exceeding the levels of the closely related serine protease-like proteins (Spls). Overall, this study provides an overview of the functional potential of the genomic landscape of JSNZ and constitutes a valuable resource for additional, detailed studies of the community.

## 2. Material and Methods

### 2.1 Bacterial strain and considered genome sequences

*S. aureus* strain JSNZ originates from preputial gland abscesses of C57BL/6 mice observed in the animal facility of the University of Auckland (Holtfreter *et al*., 2013). Glycerol stocks of JSNZ were stored at - 80 °C and thawed for 2 min at 37 °C prior to every cultivation. Annotated genome sequences of additional strains were downloaded from NCBI (accession of *Aureo*Wiki strains see provided information on https://aureowiki.med.uni-greifswald.de/Help:Contents; CC88 strains: muCC88d (GCF_003717375.1), AUS0325 (NZ_LT615218), 21343 (GCF_000245595.1)). Since there is no publicly available genome sequence for WU1, only information given in the publication could be considered (Sun *et al*., 2018). The CC88 strains were assigned to *spa* types t729 (JSNZ), t786 (21343), t7321 (AUS0325), t2311 (muCC88d), and t186 (WU1), respectively (http://spatyper.fortinbras.us/). Agr typing was performed based on the reference sequences of agrC (AF210055.1, AF001782.1, AF001783.1, AF288215.1). Whole genome-based taxonomic analysis of the *Aureo*Wiki strains, including JSNZ, was performed on the Type (Strain) Genome Server (https://tygs.dsmz.de; (Meier-Kolthoff *et al*., 2022; Meier-Kolthoff and Göker, 2019) and on PubMLST (https://pubmlst.org/; (Jolley *et al*., 2018).

### 2.2 Isolation of genomic DNA

To isolate genomic DNA for whole genome sequencing, *S. aureus* JSNZ was cultivated in 60 mL tryptic soy broth (TSB) at 37 °C with orbital shaking at 220 rpm until OD_540nm_ ≈ 1.3. An equivalent of 40 OD units of the bacterial culture was harvested with 3 mM EDTA by centrifugation for 5 min at 5,000 xg. The pellet was rinsed twice with 1 mL TE buffer (10 mM TRIS-HCI, 1 mM EDTA, pH 8.0) and centrifugation for 3 min at 16,000 xg, before being resuspended in 250 µL 10 mM TRIS-HCl pH 8.0. The suspension was immediately incubated with 20 µL lysostaphin (stock solution 5 mg/mL) for 30 min at 37 °C. To remove RNA and protein contamination, the sample was first incubated with 15 µL MgCl_2_ (200 mM) and 1 µL of each of RNaseA (2 U/µL), RNase I (20 U/µL) and RNase III (1 U/µL) for additional 15 min at 37 °C followed by incubation with 8 µL CaCl_2_ (200 mM) and 5 µL Proteinase K for another 30 min at 37 °C. Samples were mixed by flicking before every incubation step. After incubation, 100 µL 10% (w/v) SDS were added and samples gently shaken by hand until cleared up. Separation of DNA from debris was mediated by addition of 100 µL Sodium-Perchlorate (5 M) and 600 µL of Chloroform/lsoamylalcohol (24:1) followed by 30 s shaking by hand and centrifugation for 10 min at 16,000 xg. Approx. 300 µL of the upper phase were transferred into a new tube and filled up with 800 µL ice-cold 100% (v/v) ethanol (EtOH). The tube was inverted until the precipitated DNA became visible. The DNA was again carefully transferred to a new tube already containing 1 mL 70% (v/v) EtOH using a cut pipette tip. After centrifugation for 1 min at 16,000 xg, the supernatant was removed and the DNA pellet was dissolved in 300 µL nuclease-free water by flicking. To evaporate residual EtOH, the DNA sample was first incubated for 5 min at 65 °C with open lid, followed by a further 40 min at 42 °C with closed lid to ensure complete dissolving of the DNA. The DNA concentration was determined from appropriate sample dilutions with the Qubit™ dsDNA BR Assaykit on a Qubit™ 3 Fluorometer (Invitrogen™).

To deplete small fragments of genomic DNA, we performed size selection of 5 µg DNA with a PEG8000 solution (9% (w/v) PEG8000, 1 M NaCl, 10 mM TRIS-HCl, pH 8.0 in nuclease-free water). Sample volume was adjusted to 60 µL, mixed by flicking with 60 µL PEG8000 solution and centrifuged for 30 min at 10,000 xg. After removing the supernatant, the pellet was washed twice with 200 µL 70% (v/v) EtOH and centrifugation for 2 min at 10,000 xg. The pellet was air dried shortly and dissolved overnight at 4 °C in 50 µL low TE buffer (10 mM TRIS-HCI, 0.1 mM EDTA, pH 8.0). Sample integrity and successful size selection was confirmed on a 4150 TapeStation system (Agilent). All DNA samples were stored at -20 °C.

### 2.3 Whole genome sequencing and genome assembly

The circular genome of JSNZ was assembled by combining long and short sequencing reads. Short-read sequencing was performed *via* NovaSeq Illumina technology (2×150) provided by Genewiz® (Acenta Life Sciences). Long-read sequences were generated in-house on a MinION Mk1B sequencer (Oxford Nanopore Technologies). Therefore, the long-read genomic library was prepared using the Rapid Barcoding Kit 24 V14 (SQK-RBK114.24; Oxford Nanopore Technologies). Sequencing took place in a FLO-MIN114 flow cell with MinKNOW software (v24.02.16; Oxford Nanopore Technologies). Base calling for MinION reads was performed with Dorado (v7.3.11).

For genome assembly, the raw reads were quality and barcode trimmed. Illumina reads were trimmed using Trimmomatic (v0.39; Bolger *et al*., 2014). MinION reads were trimmed using Filtlong (v0.2.1; Wick *et al*., 2023), keeping the best 90% of reads longer than 1,000 bp based on the trimmed Illumina reads as external quality reference. Quality of raw and trimmed reads was assessed by FastQC (v0.11.9; https://www.bioinformatics.babraham.ac.uk/projects/fastqc/) and seqstats (https://github.com/clwgg/seqstats; Table S1). Subsequently, trimmed long reads were assembled using the Trycycler pipeline (v0.5.4; Wick *et al*., 2021). Here, reads were sub-sampled in 16 sets. Four of the subsets were each assembled using the long read assembler Flye (v2.9.3; Kolmogorov *et al*., 2019), Minipolish+Miniasm (v0.1.3 Wick and Holt, 2021) using Miniasm (v0.3; Li, 2016), Raven (v1.8.3; Vaser and Šikić, 2021) and Canu (v2.3; Koren *et al*., 2017). Draft assemblies were manually checked and curated for structural quality using Bandage (v0.8.1; Wick *et al*., 2015). Contigs of draft assemblies were clustered and resulting clusters were manually checked for outliers using FigTree (v1.4.4; http://tree.bio.ed.ac.uk/software/figtree/). The cluster representing the complete bacterial chromosome was used for reconcilation of the draft assemblies and constructing a consensus assembly based on the ONT reads. The consensus assembly was polished twice with the ONT reads using Racon (v1.5.0; Vaser *et al*., 2017) with Minimap2 (v2.26; Li, 2018) and twice with the Illumina reads using PolyPolish (v0.6.0; Wick and Holt, 2022) with bwa mem (v0.7.17; Li, 2013).

Quality of the final assembly was assessed and validated manually by comparing it to the former JSNZ assembly (GCF_003717315.1) as well as to assemblies of derivative strains constructed in parallel *via* Mauve (Darling *et al*., 2004) incorporated in Geneious Prime (v2023.0.1; Biomatters). Critical regions (e.g., SNVs, *jep* region) were re-sequenced *via* Sanger sequencing (Eurofins Genomics).

### 2.4 Integration into *Aureo*Wiki and functional annotation

Genome annotation was performed using PGAP (v2024-07-18.build7555; Li *et al*., 2021). Annotated pseudogenes were re-evaluated manually and the annotation was adjusted if necessary (e.g. separate *hisIE* and *hisF*). Regulator binding sites in the JSNZ genome were predicted based on consensus motifs using MAST (v5.3.3; (Bailey and Gribskov, 1998)) considering 300 bp upstream and 100 bp downstream of gene start codons with a e-value threshold of 1. Regions around the start codon were extracted using bedtools getfasta (v2.27.1; (Quinlan and Hall, 2010)). Motifs were generated by MEME (v5.3.3; (Bailey and Elkan, 1994) using known Staphylococcocae binding site sequences provided by RegPrecise (Novichkov *et al*., 2013) as well as the *S. aureus* WalR and GraR binding sites as used in Mäder *et al*. (2016) and SarA binding sites as described in Oriol *et al*., (2021). To obtain regulons based on the predicted binding sites, operon structures were taken into account as predicted by Operon-mapper (Taboada *et al*., 2018). In addition, regulons of *S. aureus* strains NCTC8325 (CC8) and N315 (CC5) as provided by *Aureo*Wiki were mapped to the JSNZ genome using the pan genome information provided in *Aureo*Wiki. The newly predicted and mapped regulon information was joined to gain comprehensive regulon information for JSNZ.

This curated version of the JSNZ genome was made available *via Aureo*Wiki (https://aureowiki.med.uni-greifswald.de) by integration into the already existing *S. aureus* pangenome (Fuchs *et al*., 2018), while retaining the current panIDs. For this purpose, first, a new temporary pangenome was created using the software tool PanGee, which includes the 33 strains already represented in *Aureo*Wiki and JSNZ. Based on this, each JSNZ gene was assigned to the corresponding pangene of the existing *Aureo*Wiki pangenome, such that the pangene with the most common orthologs in the new temporary and the currently existing *Aureo*Wiki pangenomes was selected. The *Aureo*Wiki integration pipeline furthermore encompassed the collection and prediction of gene and protein metadata - covering aspects such as subcellular localisation of the encoded proteins and their functional classification into families and roles (Fuchs *et al*., 2018 and “Getting Started” page of *Aureo*Wiki). For downstream analyses, data on protein localisation predicted by DeepLocPro (Moreno *et al*., 2024) and functional role assignments based on TIGRFAMs protein families (Haft *et al*., 2013) were adopted from *Aureo*Wiki.

In addition to the functional annotations introduced during *Aureo*Wiki integration, gene classifications were achieved through orthologue-based mapping to external datasets. This included, firstly, the definition of essential genes based on the study by Chaudhuri *et al*. (2009) in *S. aureus* strain SH1000 (NCTC 8325 derivate), and secondly, the association of JSNZ genes with iModulons defined by Poudel *et al*. (2022) in USA300 LAC, using the “*S. aureus* Precise165” dataset accessible *via* iModulonDBc (Catoiu *et al*., 2025). Although iModulons are inferred from transcriptomic data, they enable deeper interpretation by organising genes into conditionally co-expressed modules, which can reflect regulatory influences not captured by traditional factor-based annotations.

### 2.5 Analysis of orthologous genes and genome structure

An orthologue table containing all incorporated *S. aureus* strains is available at *Aureo*Wiki. Orthologous genes between the CC88 strains considered were determined using roary (v3.13.0; Page *et al*., 2015). Sequence similarity of orthologous genes was calculated using pairwise alignment scores provided by the R package Biostrings (Pagès *et al*., 2017), missing orthologs were replaced by a placeholder sequence containing a single ‘A’. To normalise for sequence length and composition, alignment scores were divided by the square root of the product of the respective self-alignment scores. Overall orthologous sequence similarity between groups was assessed using the median of the corresponding alignment scores.

Genome-wide comparison of orthologous gene presence was visualised using the R packages ComplexHeatmap (v2.22.0; Gu *et al*., 2016) and circlize (v0.4.16; Gu *et al*., 2014), respectively. In addition, structural comparisons of selected genomes and genomic regions were performed using NCBI BLASTn (Camacho *et al*., 2009) and the Artemis Comparison Tool (v18.2.0; Carver *et al*., 2005) as well as Lovis4u (v0.1.5; (Egorov and Atkinson, 2025). Individual orthologs of interest were manually inspected in Geneious Prime using integrated sequence Geneious Alignment and Geneious Tree Builder algorithms. Sequence alignments were visualised using the R package seqvisr (v0.2.7; (Raghavan, 2022)), and specific protein domains were annotated based on results obtained from web based InterPro searches (Blum *et al*., 2025; Jones *et al*., 2014).

### 2.6 Identification of genomic islands

The extent and type of the common staphylococcal genomic islands νSaα, νSaβ and, νSaγ within the JSNZ genome were delineated based on literature references (Baba *et al*., 2008; Gill *et al*., 2005; Kläui *et al*., 2019). Similarly, the prophage φLabRodCC88_3 was annotated according to its previous description in the literature (Yebra *et al*., 2024). Further potential genomic islands were identified based on web resources IslandViewer4 (Bertelli *et al*., 2017), PHASTEST (Wishart *et al*., 2023) and PAIDB (v2.0; (Yoon *et al*., 2015). Absence of a SCCmec was verified using SCCmecFinder (v1.2; Kaya *et al*., 2018). To classify prophages detected in the CC88 genomes, homologous prophages were identified from the NCBI Virus database (https://www.ncbi.nlm.nih.gov/labs/virus/vssi/). Integrase sequences were subsequently aligned to those of reference phages listed in Goerke *et al*. (2009) (see Table S2).

### 2.7 Genetic analysis of restriction modification systems

To explore the genetic basis of JSNZ’s previously reported high transformation efficiency, a detailed analysis of the strain’s restriction–modification (RM) systems was undertaken. Focus was placed on the *Sau1* system (*hsdRMS*). The target recognition domains (TRDs) of both HsdS alleles were aligned with the reference TRDs curated by Cooper *et al*. (2017), enabling prediction of JSNZ’s specific recognition and methylation motifs. These were derived from the best sequence matches and comparative data available for the human-associated CC88 strain AUS0325 (Kpeli *et al*., 2017). To assess sequence variability in HsdS and HsdR between the analysed *S. aureus* strains, protein sequences were aligned as described above.

Furthermore, genes JSNZ_000027 to JSNZ_000029 were identified as putative components of a type IV restriction system through BLASTp analysis (Camacho *et al*., 2009). This classification was validated and further refined by comparative alignments with reference sequences provided by Bell *et al*. (2024), and by domain annotation using InterProScan (Blum *et al*., 2025).

### 2.8 Sampling for proteomic analysis

For proteomic sample preparation, JSNZ was cultivated in an initial overnight preculture using a dilution series in 5 mL TSB at 37 °C with orbital shaking at 220 rpm. The following morning, a culture in mid-exponential growth phase (OD₅₄₀_nm_ 0.8 - 1.0) was selected to inoculate a second preculture in either TSB or RPMI medium, cultivated under the same conditions. Finally, main cultures were inoculated from the respective exponentially growing preculture (TSB: OD₅₄₀_nm_ 1.0; RPMI: OD₅₄₀_nm_ 0.5) to a starting OD₅₄₀_nm_ of 0.05 in four bioreplicates. Cultivation was carried out in a water bath at 37 °C under linear shaking at 130 rpm. Samples were taken at exponential growth phase (OD₅₄₀_nm_ 1.0 and 0.5, respectively) and 3 h after entering the stationary phase (approx. OD₅₄₀_nm_ 16.0 and 1.3, respectively).

For the cellular proteome, an equivalent of 7 OD units of bacterial culture each was rapidly cooled in liquid nitrogen (LN2) for several seconds and pelleted by centrifugation in a swing-bucket rotor for 10 min at 4816 xg and 4 °C. The supernatant was decanted, the pellets resuspended in the remaining liquid, transferred to a 1.7 mL tube and centrifuged again (1 min, 10,000 xg, 4 °C). After removing the supernatant, the pellets were washed with sterile PBS and frozen in LN2. Samples were stored at - 70°C.

For the exoproteome, 15 mL (TSB) or 35 mL (RPMI) of the culture were cooled and centrifuged as described for the cellular samples. The culture supernatant was then transferred to new tubes, leaving 1-2 mL back in the tube to prevent transfer of bacterial cells from the pellet in the supernatant samples. The supernatant samples were frozen directly in LN2. Samples were stored at -70°C.

### 2.9 Proteomic sample preparation

Cell pellets were resuspended in 50 µL 20 mM HEPES buffer (pH 8.0, 1% SDS) and disrupted in a 4.8 mL Teflon vessel filled with LN2 and an 8 mm steel ball using a ball mill (3 min, 2600 rpm). The resulting powder was resuspended in 200 µL pre-heated 20 mM HEPES buffer (50 °C, pH 8.0, 1% SDS), transferred to 1.7 mL low-bind tubes, and heated for 1 min at 95 °C while shaking at 1,400 rpm. To degrade nucleic acids, lysates were cooled to room temperature, supplemented with 2 µL 1 M MgCl₂ (final 4 mM) and 1 µL benzonase (final 0.005 U/µL) and incubated for 15 min at 37 °C while shaking at 1400 rpm, followed by sonication in an ultrasonic bath for 5 min. Finally, lysates were centrifuged for 30 min at 17,000 xg and transferred to fresh low-bind tubes. Protein concentrations were quantified using the Micro BCA™ Protein-Assay-Kit (Thermo Fisher Scientific) measured with a Synergy H1 plate reader on a Take3 microvolume plate (BioTek). For mass spectrometry (MS), 4.5 µg of protein lysates were each mixed with 0.5 µg ^15^N isotope-labelled *Bacillus subtilis* protein standard (Busch *et al*., 2025) and processed *via* SP3 sample preparation as described by Reder *et al*. (2024).

From the supernatant samples, 1.2 mL (TSB) or 4.25 mL (RPMI) were precipitated in low-bind tubes with 15% (v/v) trichloroacetic acid (TCA) for 48 h at 4 °C. Precipitated proteins were pelleted by centrifugation for 10 min at 17,000 xg and 4 °C and then washed repeatedly with 750 µL ice-cold 70% (v/v) EtOH until the pellets appeared completely white. These exoproteome pellets were subsequently rinsed with 500 µL 100% EtOH, centrifuged for 10 min at 17,000 xg and 4 °C, and air-dried under a fume hood for 30 min. Dried pellets were resuspended in 20 mM HEPES buffer (pH 8.0, 1% SDS), sonicated for 2 min, and dissolved by agitation at 60 °C and 1,400 rpm for 10 min. Protein concentrations of the precipitated samples were quantified as for the cellular samples. Protein concentrations in the raw supernatants were inferred based on these quantifications and mean protein concentration was calculated per medium. Same volumes of the raw exoproteome samples as precipitated before (1.2 mL and 4.25 mL, respectively) were supplemented with the ^15^N isotope-labelled *B. subtilis* protein standard to a proportion of 20% (w/w) of the corresponding mean protein content per culture medium and precipitated as described above. For MS, 2 µg of each precipitated standard supplemented exoproteome sample was processed *via* SP3 sample preparation.

To visualise the composition of stationary JSNZ exoproteomes, a silver stained SDS-PAGE was used. Specifically, 280 µL of each stationary RPMI culture supernatant (∼ 1.1 µg protein) was lyophilised and resuspended in 9 µL of nuclease-free water. These concentrated samples, along with 9 µL of each stationary TSB culture supernatant (∼ 1.3 µg protein), were subjected to electrophoretic separation on a 4–12% Bis-Tris NuPAGE™ gel (Thermo Fisher Scientific). Silver staining was subsequently carried out as described in (Wolfgramm *et al*., 2025).

### 2.10 Mass spectrometry (MS) and data analysis

Proteomic samples were analysed *via* liquid chromatography-tandem mass spectrometry (LC-MS/MS) in data independent mode (DIA). The instrument setup consisted of an UltiMate™ 3000 RSLCnano HPLC coupled to an Orbitrap Exploris™ 480 mass spectrometer (both Thermo Fisher Scientific). For more information, please refer to Tables S3 and S4.

MS data were analysed with the Spectronaut software (v19.9.250512.62635, Biognosys) using the library-based search mode. The respective spectral libraries were composed on the basis of 102 MS measurements of JSNZ cellular and supernatant samples grown in different media. The spectra were searched according to the newly provided database of JSNZ proteins including protein sequences of annotated pseudogenes generated from the updated genome assembly. The analyses were performed with median cross-run normalisation of ion intensities on the ^15^N labelled standard (100% percentile). The labelled standard was covered by a second spectral library (Busch *et al*., 2025). The precursor q-value cut-off was set to 0.001. The spectral libraries and MS analysis output data have been deposited to the ProteomeXchange Consortium *via* the PRIDE (Perez-Riverol *et al*., 2025) partner repository with the dataset identifier PXD067993.

Intensity-based absolute quantification (iBAQ) values (Schwanhäusser *et al*., 2011) and ion-intensity values reported by Spectronaut, which had been normalised using the ^15^N-labeled internal standard, were further normalised to OD for exoproteome samples. This improved comparability, particularly between supernatant samples harvested during exponential and stationary growth phases. However, a direct comparison is only valid among all pellet samples (independent of growth medium) or among supernatants originated from the same medium. This is due to large differences in absolute protein concentrations across the different supernatants, which necessitated the use of varying amounts of internal standard. For comparisons between supernatants from RPMI and TSB media, the data were normalised using a median-to-median approach.

The normalised data were used to infer protein abundances (iBAQ) of proteins with at least two identified peptides and perform statistical analysis based on the peptide-centric ROPECA method (Suomi and Elo, 2017) implemented in the R package SpectroPipeR (v0.3.0; Michalik *et al*., 2025). The statistical analysis of abundance changes in specific proteins or protein sets was conducted using the R package rstatix (v0.7.2; (Kassambara, 2023a). Gene set enrichment analyses (Subramanian *et al*., 2005) were carried out based on the signed Euclidean distance to the zero point considering the log2 ratios and adjusted p-values of unique proteins using the R package fgsea (v1.32.4; Korotkevich *et al*., 2016).

### 2.11 Data handling and visualisation

All data were processed using R (R Core Team, 2024) *via* the graphical interface of RStudio (RStudio Team, 2020). For tabular data R packages tidyverse (v2.0.0; Wickham *et al*., 2019), readxl (v1.4.5; Wickham and Bryan, 2025), and openxlsx (v4.2.8; Schauberger and Walker, 2025) were used, FASTA files were imported and arranged using microseq (v2.1.6; Snipen and Liland, 2023).

Unless otherwise stated, data visualisation was likewise carried out in R, primarily using the ggplot2 package (Wickham, 2016) within the tidyverse framework. Its functionality was expanded through R packages scales (v1.4.0; Wickham *et al*., 2025), ggtext (v0.1.2; Wilke and Wiernik, 2022), ggrepel (v0.9.6; Slowikowski, 2024), ggpubr (v0.6.0; Kassambara, 2023b), and ggh4x (v0.3.1; van den Brand, 2025). Genomic regions mapped to corresponding protein abundances were visualised using gggenes (v0.5.1; Wilkins, 2023). Set intersection were depicted *via* R packages UpSetR (v1.4.0; Gehlenborg, 2019) and eulerr (v7.0.2; Larsson, 2024), respectively. Plot arrangements were handled using patchwork (v1.3.0; Pedersen, 2024) and gridExtra (v2.3; Auguie, 2017), colour palettes were accessed *via* paletteer (v1.6.0; Hvitfeldt, 2021).

## 3. Results

### 3.1 JSNZ reference genome

The completed genome of JSNZ comprises a circular chromosome of 2,719,558 bp with a GC content of 32,9%, 2645 annotated genes and 56 annotated pseudogenes (Supplementary data 1: Table 1-1). No plasmids were detected. Compared to the most complete JSNZ genome to date (NCBI accession GCF_003717315.1, Holtfreter *et al*., 2013), gaps of 26,848 bp in total have been closed and 80 genes have been newly defined or modified, including for example *eap* (extracellular adherence protein). The assembly level could thus be improved from a scaffold comprising 15 contigs to a single complete genome with an increase in CheckM completeness from 98.53% to 99.8%. In addition, it was integrated into the *S. aureu*s data base *Aureo*Wiki (https://aureowiki.med.uni-greifswald.de), where it is presented with comprehensive information about each gene including the predicted regulon (Supplementary data 1: Table 1-2). JSNZ represents the first mouse-adapted strain and the first CC88 strain among the now 34 *S. aureus* strains included in *Aureo*Wiki (Fig. 1A).

**Figure 1.**
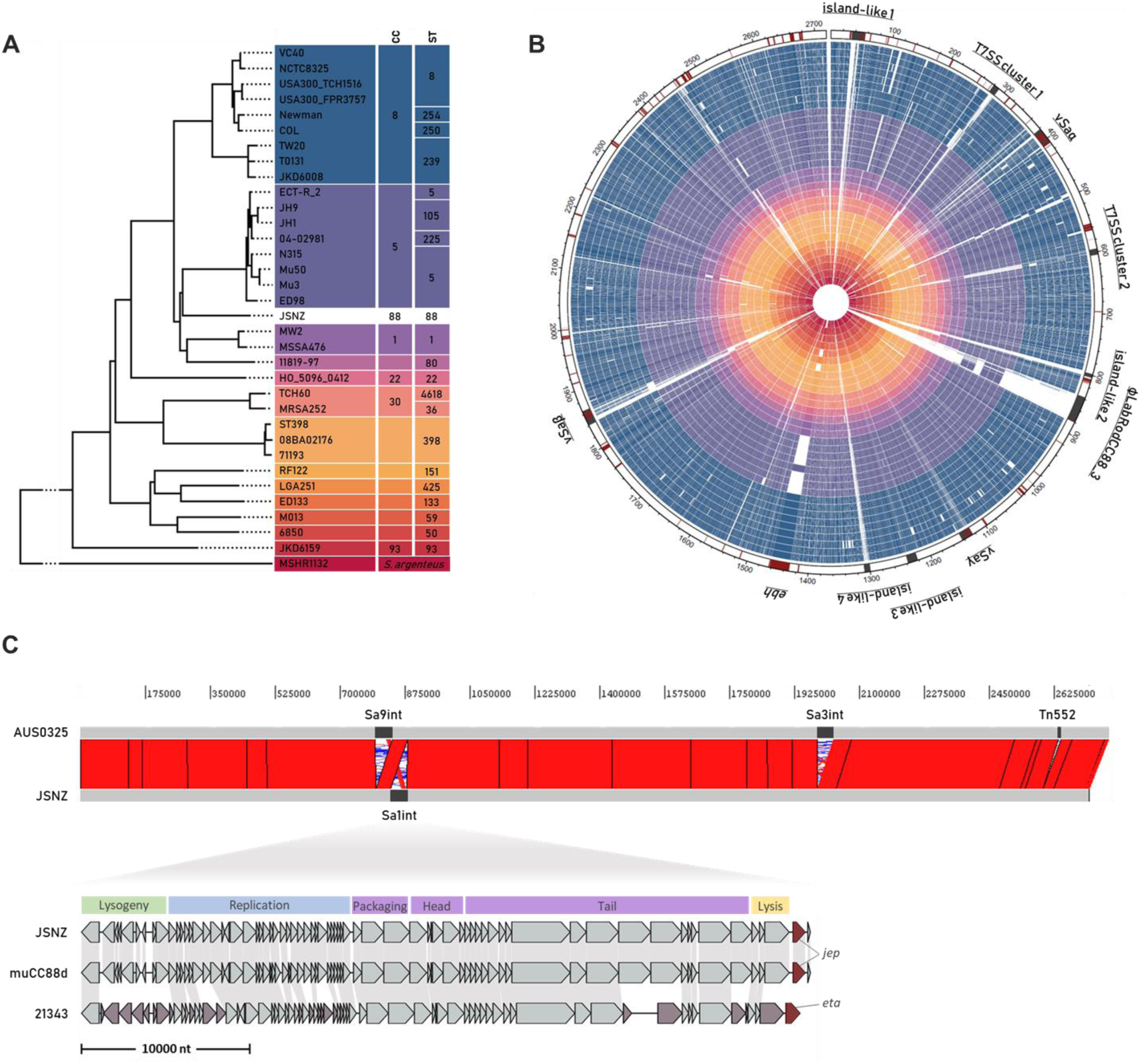
Comparison of *S. aureus* JSNZ to existing *Aureo*Wiki strains and JSNZ genome architecture. **(A)** Phylogenetic tree of all 34 strains integrated in *Aureo*Wiki indicating clonal cluster (CC) and sequence type (ST). Coloured according to CC. **(B)** Circular representation of the JSNZ genome (outer circle) and orthologous genes within the genomes of the *Aureo*Wiki strains (circles ordered and coloured like in (A)). In the outer circle, JSNZ genes associated with virulence or resistance are depicted in dark red and genomic islands and important genome regions are depicted in dark grey. Region names and genome position in kilobases indicated outside the circle. **(C)** Comparison of the genome architecture between JSNZ and the human CC88 isolate AUS0325, with a detailed view of the Sa1int phage ΦLabRodCC88_3 compared to that of further CC88 strains (incl. ΦETA2 of isolate 21343). Genome position in base pairs is indicated above. Ribbons represent sequence homology. Phage modules of Sa1int phages are provided above the detailed comparison. Phage-encoded virulence-associated genes are highlighted in dark red, genes without homologs are coloured in reddish grey.

**Table 1.**
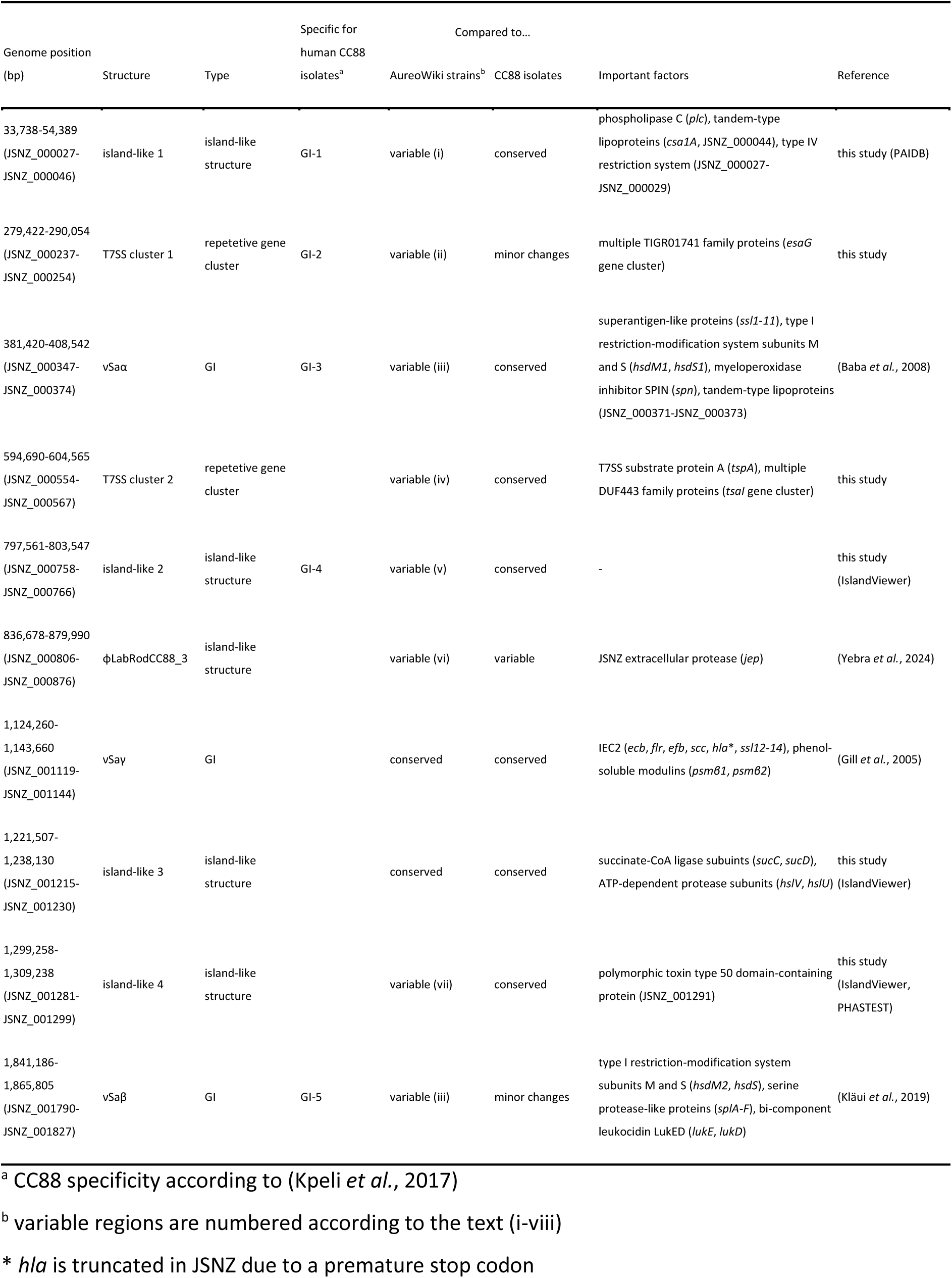
Genomic islands, island-like structures and repetitive gene clusters in the JSNZ genome.

Many of the genotypic characteristics of JSNZ have been described (Holtfreter *et al*., 2013; Schulz *et al*., 2017; Trübe *et al*., 2019) and are only briefly summarised here: JSNZ carries the type 8 capsular polysaccharides (*cap8*), the accessory gene regulator of type III (*agr3*) and a specific virulence and resistance pattern, that is mainly shaped by genomic islands (GI)(Fig. 1B, Supplementary data 2). In total, ten regions of the JSNZ genome are classified as GIs, island-like structures or repetitive gene clusters, including the well-known νSaα, νSaβ and νSaγ as well as a Sa1int phage (Table 1). Eight of these ten regions differ substantially from that of the other *Aureo*Wiki strains, i.e. show few orthologous genes in these strains (Fig. 1B, Supplementary data 4): (i) An island structure downstream of *rmlH*/*orfX* (island-like 1), which encodes two tandem-type lipoproteins (*lpl*) and a potential restriction modification system (JSNZ_000027-JSNZ_000029) that is unique among the *Aureo*Wiki strains. The methicillin resistance determining staphylococcal cassette chromosome SCCmec, which is integrated at this position in many human isolates, is not present in JSNZ. (ii) A region downstream of the type VII secretion system (T7SS) gene cluster (T7SS cluster 1), harbouring T7SS effectors as well as a repetitive cluster of TIGR01741 family proteins (*esaG* immunity gene cluster). (iii) The νSaα containing a cluster of 11 superantigen-like proteins (*ssl*), a further cluster of three tandem-type lipoproteins (*lpl*) and subunits M and S of the type I restriction-modification system (*hsdMS*). The JSNZ νSaα is comparable to type I according to Baba *et al*. (2008). (iv) The TspA region (T7SS cluster 2) encoding a T7SS effector (*tspA*) and a linked range of DUF443 family proteins. (v) An island structure upstream of the encoded virulence factors clumping factor A (*clfA*), von Willebrand factor binding protein (*vwb*) and extracellular matrix protein-binding adhesin (*emp*)(island-like 2). (vi) The Sa1int prophage φLabRodCC88_3 encoding a secreted serine protease called JSNZ extracellular protease (*jep*). (vii) An island structure (island-like 4) encoding a toxin domain-containing protein besides several hypothetical proteins. (viii) The νSaβ containing eight different serine protease-like proteins (*spl*), the bi-component leukocidin LukED (*lukED*) and a second pair of modification-restriction system subunits HsdMS. The *bsa* gene cluster as well as the enterotoxin gene cluster (*egc*) is missing. Although there is a clear similarity to νSaβ type XV (Kläui *et al*., 2019; Schwendimann *et al*., 2021), the JSNZ νSaβ differs from this type at the 5’ end and cannot be classified unambiguously.

### 3.2 Comparison of the JSNZ genome to selected CC88 genomes

Five of the eight distinct regions that distinguish JSNZ from the other *Aureo*Wiki strains correspond to the regions described as specific for *S. aureus* CC88 (GI1-5 in Kpeli *et al*., 2017), namely regions (i), (ii), (iii), (v) and (viii) (Table 1, Fig. S1). For a more detailed comparison, three further CC88 isolates of different origins and characteristics were selected: AUS0325, a human methicillin-susceptible strain from Australia (CC88 reference genome), 21343, a human methicillin-resistant strain from the United States (Kpeli *et al*., 2017), and muCC88d, an isolate derived from a wild field vole (Trübe *et al*., 2019). Because of limited genomic information available for WU1, we had to exclude this strain.

The JSNZ genome resembles the genome architecture of the human CC88 isolate AUS0325 (CC88 reference genome; Kpeli *et al*., 2017), differing only with respect to four mobile genetic elements (MGEs) (Fig. 1C, Table 2). While AUS0325 carries a *hlb*-converting Sa3int phage encoding the immune evasion cluster 1 (IEC1), a Sa9int phage and a Tn552 transposon encoding the *blaZ* gene cluster (albeit presumably non-functional due to a truncated *blaR1*), JSNZ is positive for the Sa1int phage φLabRodCC88_3. Both lack a Sa2int phage as well as the already mentioned methicillin resistance determining SCCmec. The absence of MGEs common in human isolates is consistent with the finding that adaptation to the murine host is accompanied particularly by acquisition and loss of phages (Yebra *et al*., 2024). However, not only the presence of certain phages but also the gene content of particular phages is of relevance. An important difference occurs in the Sa1int phage carried by isolates JSNZ, muCC88d and 21343 (Fig. 1C). While the respective phage in the human isolate 21343 encodes for the exfoliative toxin A (*eta*) (φETA2; NCBI RefSeq: NC_008798.1), the one in the murine isolates JSNZ and muCC88d encodes for the JSNZ extracellular protease (*jep*) at the same position (φLabRodCC88_3; Yebra *et al*., 2024). This is particularly interesting as *jep* occurs almost exclusively in mouse-derived *S. aureus* isolates (Yebra *et al*., 2024).

**Table 2.** Presence and type of mobile genetic elements (MGE) in the four CC88 genomes considered in this study. νSaα typing based on Baba *et al*. (2008), νSaβ typing based on Kläui *et al*. (2019).

| MGE | JSNZ | muCC88d | AUS0325 | 21343 |
| --- | --- | --- | --- | --- |
| phages |  |  |  |  |
| Sa1int | ϕLabRodCC88_3 (*jep*) | ϕLabRodCC88_3 (*jep*) | - | ϕETA2 (*eta*) |
| Sa2int | - | Δ*pvl* | - | - |
| Sa3int | - | *sak, scn, chp* | *sak, scn* | *sak, scn, chp* |
| Sa9int | - | + | + | + |
| GI |  |  |  |  |
| vSaα | type I | type I | type I | type I |
| vSaβ | type XV<sup>a</sup> | type XV<sup>a</sup> | type XV<sup>a</sup> | type XV<sup>a</sup> |
| vSay | *hla* * | *hla*<sup>b</sup> | *hla* | *hla* |
| resistance |  |  |  |  |
| SCCmec | - | - | - | + |
| blaZ cluster | - | - | Tn552 (*blaR1* *) | plasmid (co-carriage<br>*cadD*) |

+ presence of the element, - absence of the element, \* truncated due to premature stop codon,
<sup>a</sup> modified type XV with differing 5' genes, <sup>b</sup> muCC88d exhibits a Leu172Pro substitution, inactivating hemolysis (Trübe *et al.*, 2019).

Besides phage-located loci, only a limited number of loci is specifically enriched in *S. aureus* isolates from mice compared to those from other hosts across all CCs (Yebra *et al*., 2024). Of the 15 mouse-specific, non-phage loci described, eight appear in the JSNZ genome (Supplementary data 3: Table 3-1), including variants of the staphylococcal superantigen-like proteins 4 and 7 (*ssl4*, *ssl7*).

Further loci with potential implications for virulence and host adaptation were identified through analysis of the pseudogenes annotated in JSNZ. While some genes were truncated in JSNZ only (e.g. *ktrB*, formate/nitrite transporter JSNZ_000258) or truncated in all CC88 strains analysed in this study (e.g. *tcyP*, *msrA3*), there are also genes that were exclusively affected in the rodent isolates JSNZ and muCC88d (e.g. *hla*, *ebh*, *rbf*). As described previously, both murine isolates lack hemolytic activity of alpha-hemolysin (Hla) as they carry mutated *hla* versions, leading to a truncated Hla in JSNZ and a Hla Leu172Pro substitution in muCC88d (Trübe *et al*., 2019). Similarly, the regulator of biofilm formation (Rbf) – a helix-turn-helix transcriptional regulator (Lim *et al*., 2004) – is truncated by different mutations in JSNZ (frameshift) and muCC88d (substitution). In case of the giant ECM-binding protein homologue (Ebh) both murine CC88 strains exhibit the same premature stop codon and a similar insertion of 79 amino acids that is unique among the strains considered (Fig. 2B). This coincidence is noteworthy given that JSNZ and muCC88d are of different origin (laboratory-mice and wild rodent, respectively, Trübe *et al*., 2019).

**Figure 2.**
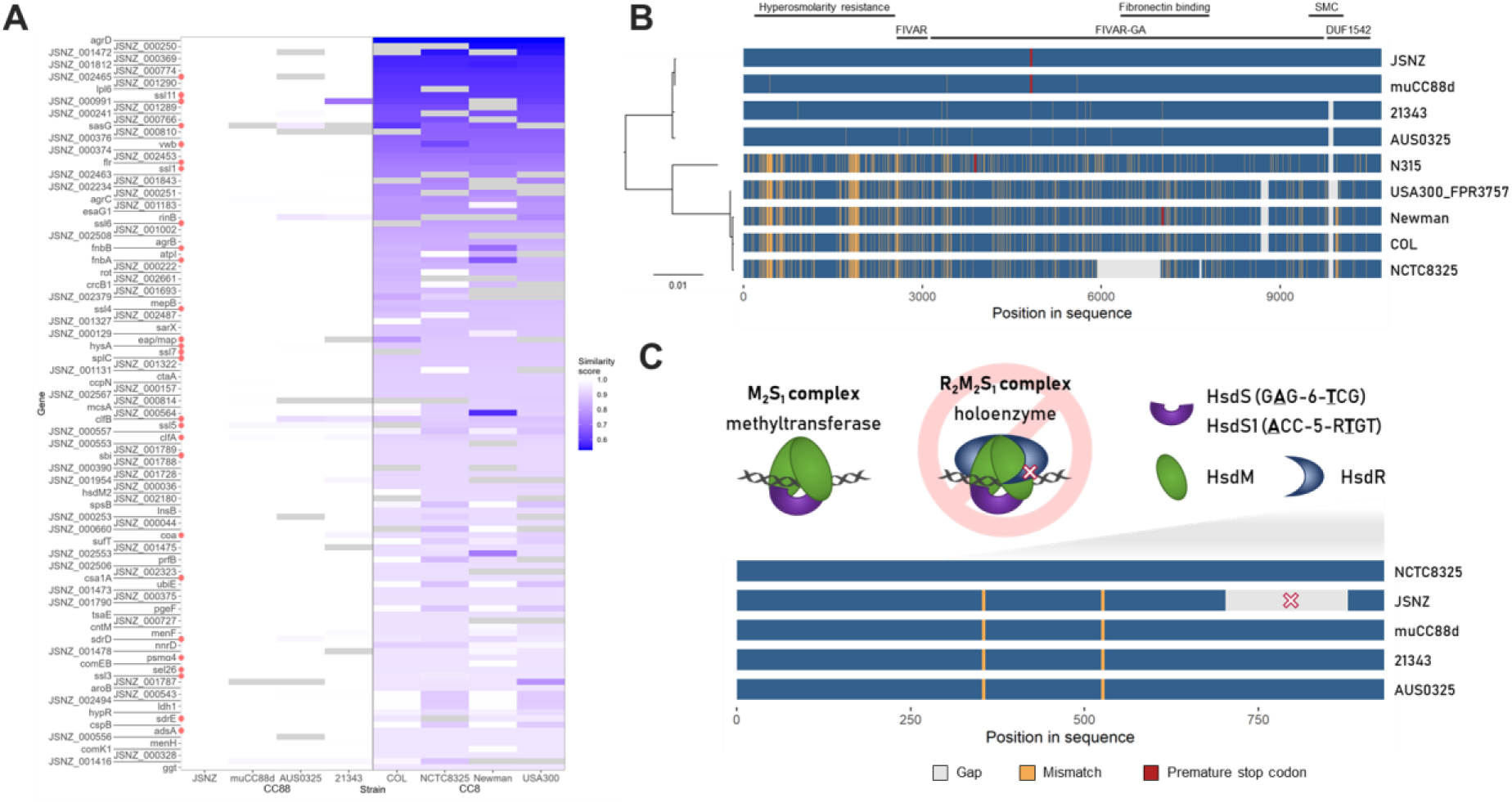
Sequence similarity of orthologous genes of CC8 reference strains and selected CC88 isolates. **(A)** Protein sequence similarities of genes with highest deviation between CC8 and CC88 variants (similarity < 95%). JSNZ variant used as reference, grey fields reference to missing orthologs. Virulence associated genes are labelled by red dots. **(B)** Protein sequence alignment of ECM-binding protein homologue (Ebh) orthologs. Amino acid mismatches (yellow), gaps (grey) and premature stop codons (dark red) are depicted. Coverage of JSNZ Ebh by mass spectrometric measurement is indicated by green lines in the upper sequence. Known domains and domain functions are indicated above the alignment (SMC = structural maintenance of chromosomes domain). **(C)** Scheme of Type I restriction-modification system subunits and theoretically built functional complexes M_2_S_1_ (methylation) and R_2_M_2_S_1_ (restriction and methylation). Protein sequence alignment of Type I restriction endonuclease HsdR orthologs with highlighted restriction-abolishing deletion. Colouring as in (B).

### 3.3 Comparison of orthologous protein-coding genes between JSNZ and common CC8 reference strains

Several common *S. aureus* reference strains belong to CC8, and strains from this cluster are highly prevalent in clinics (Albrecht *et al*., 2015; Tabatabaie Poya *et al*., 2025). To evaluate gene-level differences between JSNZ and the most common CC8 references strains COL, NCTC8325, Newman, and USA300, the amino acid sequence similarity of their respective orthologous gene products was compared. The three selected CC88 isolates were included in the analysis to infer whether the observed differences may be lineage-specific or unique to JSNZ.

Among the 2,342 orthologs analysed, 129 gene products exhibited less than 95% amino acid sequence similarity between the compared CC8 and CC88 isolates (based on group-wise similarity score medians), including several well-known lineage-specific proteins such as Agr components, Hsd subunits, and T7SS immunity proteins (Fig. 2A, Supplementary data 1: Table 1-3). The most pronounced difference was observed for AgrD, with only 53% sequence similarity. Several virulence factors also belong to this group of differing proteins, including ClfA, ClfB, FnbA, FnbB, Coa, vWbp, HysA, SplC, Psmα4, Sel26, Sbi, AdsA, SasG, SdrD, SdrE, and FLIPr, as well as multiple Lpls and SSLs. Among the SSLs are the already mentioned SSL4 and SSL7, which occur in a mouse-associated variant in JSNZ (Yebra *et al*., 2024), and SSL11, whose JSNZ gene variant was previously identified as *ssl11_JSNZ* (Holtfreter *et al*., 2013). Notably, these three SSLs share identical amino acid sequences across the analysed CC88 isolates, including both human-derived isolates.

A high sequence similarity among the four CC88 isolates was observed for the majority of the 129 gene products differing between CC8 and CC88. This has already been described for ClfA (Sun *et al*., 2018). Only ClfB and RinB show additional variation between human and murine CC88 isolates (Fig. 2A). A rather complex pattern emerges for the surface protein SasG, which consists of an N-terminal A domain, responsible for bacterial adhesion to desquamated epithelial cells (Roche *et al*., 2003b), and a variable number of B repeats (Roche *et al*., 2003a). While the number of B repeats varies within the CCs, the A domain is largely similar between human and murine CC88 isolates, but differs between the analysed CC8 and CC88 strains (Fig. S2).

### 3.4 Restriction-modification systems in JSNZ

JSNZ was reported to be efficiently transformable (Holtfreter *et al*., 2013). Therefore, we focused on the restriction modification systems present in JSNZ. Three systems were identified, the well-known Sau1 type I restriction-modification system (Sau1RM, Hsd), the SauUSI type IV restriction enzyme (SauUSI homologue JSNZ_002468) and the potential system encoded in the island-like structure 1. The Sau1RM system is known to provide a lineage specific transformation barrier. Consistent with this, the target recognition domains (TRDs) of JSNZ’s Sau1RM specificity subunits (HsdS and HsdS1) are virtually identical to those of the other CC88 strains. Alignment with the TRD sequences published by Cooper *et al*. (2017) revealed a combination of TRDs N and Q for HsdS1 (JSNZ_000367) and TRDs NOVEL1 and K for HsdS (JSNZ_001801) leading to the recognition sequences <u>A</u>CC-5-R<u>T</u>GT and G<u>A</u>G-6-<u>T</u>CG, respectively (Fig. S3; Kpeli *et al*., 2017). However, the JSNZ Sau1RM restriction subunit HsdR exhibits a 172 aa deletion (Δ705-876) in the C-terminal domain (CTD) while the other three investigated CC88 strains carry a full-length version of HsdR (Fig. 2C).

The SauUSI homologues of the four strains only showed minor substitutions to the NCTC8325 reference (SAOUHSC_02790). The additional potential restriction system (JSNZ_000027-JSNZ_000029) is encoded on the island-like structure 1, i.e. in a specific for CC88 strains. However, in AUS0325 two of the three genes are disrupted by premature stop codons. Sequence homology revealed the system as a member of the just recently defined type IV restriction system subclass of coiled-coil nuclease tandems (CoCoNuTs), specifically a type I-B CoCoNuT (Fig. S4; Bell *et al*., 2024). This subclass is assumed to predominantly target RNA, although this remains to be definitively established.

### 3.5 Genome coverage by the proteome under laboratory conditions

Based on the annotated genome sequence, we characterised which predicted coding sequences (CDS) are actually translated into proteins under laboratory conditions. We considered standard liquid cultures in a rich medium (TSB), as well as in a minimal medium relevant for cell culture infection models (RPMI), examining both the exponential and stationary growth phase (Fig. 3A, Fig. S5, Fig. S6). Of the 2,620 predicted non-RNA genes (incl. pseudogenes), 2,050 (78.2%) were detected to be translated into a protein in at least one of these conditions, most could be detected in all four conditions (1,738 / 66.3%; Fig. 3B, Supplementary data 1: Table 1-4). This covers 89.9% of the predicted cytoplasmic proteins, 75.8% of the cytoplasmic membrane proteins, 100% of the cell wall and surface proteins, and 76.8% of the extracellular proteins (according to DeepLocPro, Fig. 3C).

**Figure 3.**
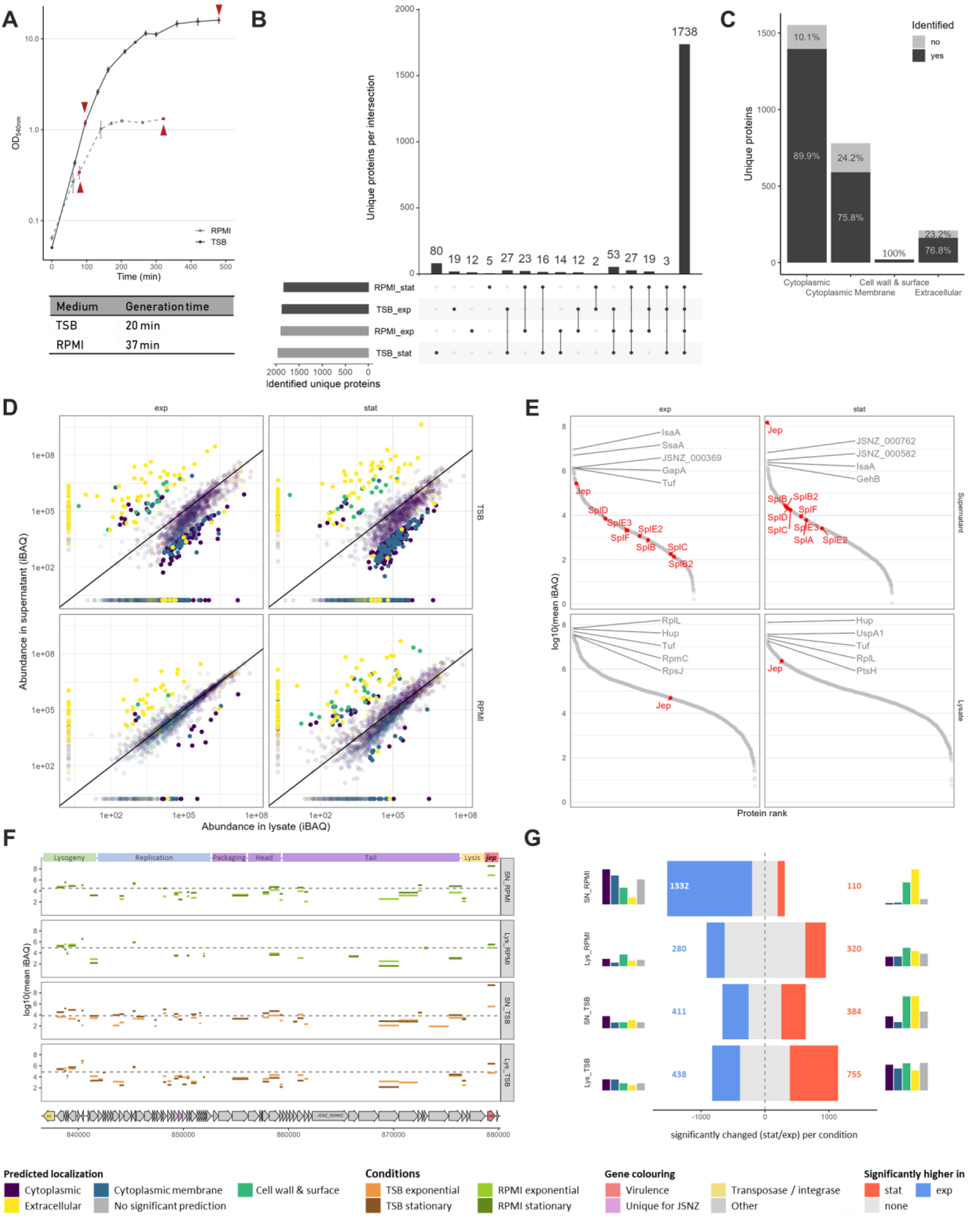
Proteome and exoproteome analysis of JSNZ grown in TSB and RPMI. **(A)** Growth curves of JSNZ cultivated in TSB (solid line) or RPMI (dashed line). Sampling time points are indicated by dark red arrowheads, generation time during exponential growth phase indicated below the plot. Error bars indicate standard deviation. **(B)** Intersections among experimental conditions based on presence of the 2,050 proteins identified via mass spectrometry (MS). Horizontal bars at the left show total number of unique proteins identified per condition; vertical bars represent the number of proteins shared across specific combinations of conditions. **(C)** Coverage of the JSNZ genome by detected proteome, represented by percentage of predicted genes for which a protein product was identified in at least one condition, grouped by predicted subcellular localisation (DeepLocPro). **(D)** Relative abundance of proteins in the cellular lysate compared to the relative abundance in the supernatant samples per condition. Abundances are inferred from median-median normalised iBAQ values. iBAQs of proteins identified in only one fraction were set to half-minimal iBAQ in the missing one. Proteins are coloured by predicted subcellular localisation, proteins with normalised iBAQ >1×10^4^ and absolute fold-change >20-fold are depicted in bright colours. **(E)** Abundances of proteins identified in the samples derived from TSB cultures (all samples including RPMI shown in Fig. S5). Secreted serine-protease like proteins (Spls) and JSNZ extracellular protease (Jep) are labelled in red, top five proteins based on abundance are labelled in grey. **(F)** Proteomic coverage of ΦLabRodCC88_3 encoded proteins depicted by protein abundance per sample. The annotated ΦLabRodCC88_3 is shown with genome position in base pairs, phage modules are indicated at the top of the plot. Median iBAQ per medium and fraction is indicated by a dashed line, protein iBAQ lines are coloured based on condition. **(G)** Number of proteins significantly altered between stationary and exponential growth phases based on ROPECA statistics (absolute fold-change >1.5; q < 0.05) across conditions. Left and right bar charts depict the proportional composition of predicted localisations of altered proteins. Colour legend for all plots summarised in this figure is given at the bottom. All values based on four bioreplicates. Lys = lysate/cellular proteome, SN = supernatant/exoproteome.

However, while the localisation predicted by DeepLocPro generally correlated with the relative abundance pattern, with cytoplasmic and membrane proteins enriched in the pellet fraction, and cell wall, surface, and extracellular proteins enriched in the supernatant, there are a few outliers (Fig. 3D). Outliers were defined as proteins with median-normalised iBAQ values ≥ 1×10⁴ that exhibited opposing abundance trends across all four conditions, and showed at least a 20-fold difference in abundance between the pellet and supernatant fractions in at least one condition. Among the outliers are proteins assigned to the Sa1int prophage φLabRodCC88_3, the potential CoCoNuT restriction system, and proteins related to T7SS (Supplementary data 3: Table 3-2). Part of the discrepancy between predicted and observed results could be - besides an erroneous prediction - due to atypical release, as some of the proteins were previously detected in extracellular membrane vesicles (LtaS, FtsL, RpmB, MsaA, JSNZ_001786, JSNZ_001787, JSNZ_001899, JSNZ_002410; EVpedia (Kim *et al*., 2015) and Uppu *et al*., 2023), or were released from the membrane by proteolytical cleavage (LtaS, JSNZ_000664, JSNZ_001322; Cheng *et al*., 2020; Nega *et al*., 2015).

A total of 42 proteins belonged to the top 100 most abundant proteins in all exoproteome samples, while 63 proteins belonged to the top 100 in all cellular proteome samples (Supplementary data 1: Table 1-5). These groups overlapped in 33 proteins that were highly abundant in both the pellet and the supernatant, with the majority encoded by essential genes as defined by Chaudhuri *et al*. (2009). The most abundant proteins in the pellets were mainly proteins involved in genome integrity and protein biosynthesis accompanied by some central factors of metabolism. In the supernatants, highly abundant proteins include factors like IsaA, Sbi, IsdA, GehB, SSL11 and Jep (Fig. 3E, Fig. S5). Jep is particularly noteworthy, as it is by far the most abundant protein in the supernatant during the stationary growth phase. In TSB medium, it accounts for an impressive 74.1% of the total protein intensity (iBAQ) of the stationary supernatant sample (RPMI: 16.5%). The rest of φLabRodCC88_3 – where Jep is encoded in the terminal part – reflects the typical pattern of a lysogenic phage. There is an increased protein abundance among the lysogeny module and a low abundance of most other phage proteins. However, the CI- and Cro-like repressors could not be detected in the proteome (Fig. 3F).

Only five of the 56 pseudogenes were found to be translated into a protein in at least one condition, three of which could be detected under all conditions examined (Hla, Ebh, tandem-type lipoprotein JSNZ_002466; Fig. S7). Interestingly and despite the premature stop codon, Hla is found in rather high amounts and the shortened ORF is well covered by identified peptides (Supplementary data 3: Table 3-3).

### 3.6 Proteome changes in response to growth phase and culture medium

The proteome of JSNZ differed significantly between the compared media and growth phases (Fig. 3G, Supplementary data 1: Tables 1-6 & 1-7). When comparing the exponential to the stationary growth phase in the TSB medium, about 60% of the detected proteins were significantly altered in their abundance (OD normalised, |fold-change| > 1.5, adjusted p-value < 0.05). This is the case for both the cellular proteome and the exoproteome. In RPMI medium, the cellular proteome changed less between growth phases, while the exoproteome showed a rather large number of altered proteins. Of note, this marked change in the exoproteome was mainly driven by proteins with predicted cytoplasmic localisation (Fig. 3G). To characterise the extensive changes between the conditions, the alterations were analysed with regard to TIGRFAM main and sub role (functional classification), assignment to iModulons according to Poudel *et al*. (2022), and predicted regulons (regulatory classifications; Supplementary data 1: Table 1-1). Although iModulons and regulons widely yield overlapping results, both classifications were used in the analysis to also capture nuanced regulatory insights. While regulons are defined mechanistically by shared transcriptional regulators, iModulons group conditionally co-expressed gene sets independently of specific regulation factors (e.g. virulence iModulons Vir-1 and Vir2; Catoiu *et al*., 2025).

When cultivated in the complex medium TSB, the proteome of JSNZ traced the change from strong proliferation to cell survival (Fig. 4A, Fig. S8). Proteins with functions in protein biosynthesis, DNA metabolism and cell envelope-associated processes were significantly more abundant during the exponential growth phase than during the stationary phase. In contrast, the stationary phase proteome was characterised by proteins involved in energy metabolism, amino acid and nucleotide metabolism as well as signal processing, i.e. various stress responses (SigB, Rex, CodY). This pattern was also reflected by the enrichment of corresponding regulons and iModulons (Fig. 4A). The enrichment analysis of these regulatory groups furthermore revealed an elevated induction of protein stress responses (CtsR, HrcA) during the exponential phase, likely linked to the high translational activity.

**Figure 4.**
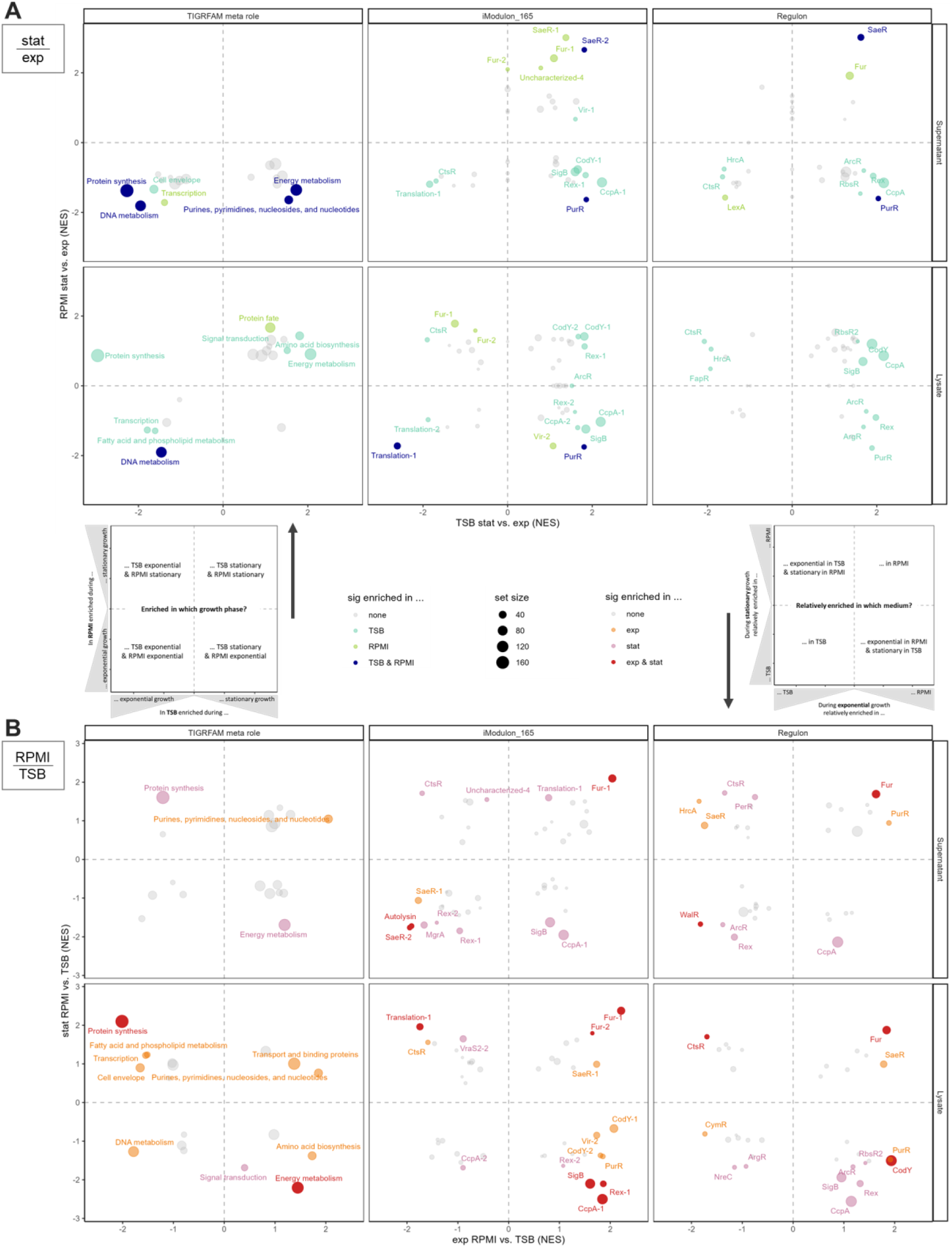
Enrichment of protein sets comparing growth phases and media. **(A)** GSEA statistics for proteome alteration from exponential to stationary growth phase per medium and **(B)** for relative proteome alteration from TSB to RPMI per growth phase. Proteins were grouped to sets according to TIGRAFM meta role (left), assigned iModulon (centre), and regulon (right), respectively. Points are coloured based on significant enrichment (adjusted p-value <0.05) in one or both respective comparisons (four bioreplicates). Point size is relative to protein set size. Legends are centrally collected. Explanatory, schematic plots are included to aid data interpretation. NES = normalised enrichment score.

Contrary to the situation in TSB, the proteome in the synthetic medium RPMI reflected a parallel manifestation of growth and survival with a less pronounced transition between the growth phases. This was accompanied by a prolonged doubling time in RPMI compared to TSB (TSB: 20 min, RPMI: 37 min; Fig. 3A). Although similar trends as in TSB were observed for the most functional categories from exponential to stationary phase in RPMI (decrease in transcription and translation, increase in energy metabolism and signal transduction), most sets were not significantly enriched (Fig. 4A). The contrast between the two media became even more apparent when comparing the relative proteome compositions (Fig. 4B). During exponential growth, proteins involved in protein synthesis were significantly enriched in TSB compared to RPMI, whereas during stationary phase, they were significantly enriched in RPMI. The opposite pattern was observed for the overall energy metabolism (TIGRFAM meta role), which showed higher enrichment in TSB during the stationary and in RPMI during exponential phase. This trend was particularly evident for the iModulons SigB, Rex-1, PurR and CcpA-1, as well as the CodY regulon. Notably, only the Fur iModulons exhibited significantly higher proportions in RPMI across both growth phases (Fig. 4B), with increasing abundance from exponential to stationary phase (Fig. 4A).

### 3.7 Expression of virulence factor genes

As JSNZ is of particular interest as reference strain for murine infection models, we specifically analysed the expression of virulence factors under the tested laboratory conditions. Of the 108 virulence factors defined in the JSNZ genome, 99 were detected in the exoproteome under at least one condition (missing: IcaABCD, Sel26, Lpl6, Psmα2, Psmα3, VraX; Table S2). The translated virulence factors were mainly assigned to immune evasion and adhesion (25% and 29%, respectively, incl. haemoglobin binding and secreted clotting factors), followed by secreted enzymes (19%), cytolytic toxins (17%) and tandem-type lipoproteins (10%; Fig. 5A). However, the intensity pattern of the respective virulence factor groups differed between TSB and RPMI (supernatant samples of stationary growth phase; Fig. 5B): While secreted enzymes and cytolytic toxins were predominant in the complex medium TSB, secreted adhesins/clotting factors and haemoglobin binding proteins were prominent in the minimal medium RPMI. Furthermore, proteins involved in immune evasion were, in general, also increased in RPMI and reached comparable levels to the other major virulence groups. Five virulence factors belonged to the top 100 proteins in all supernatant samples, namely Jep, SSL11, IsaA, IsdA, and Sbi.

**Figure 5.**
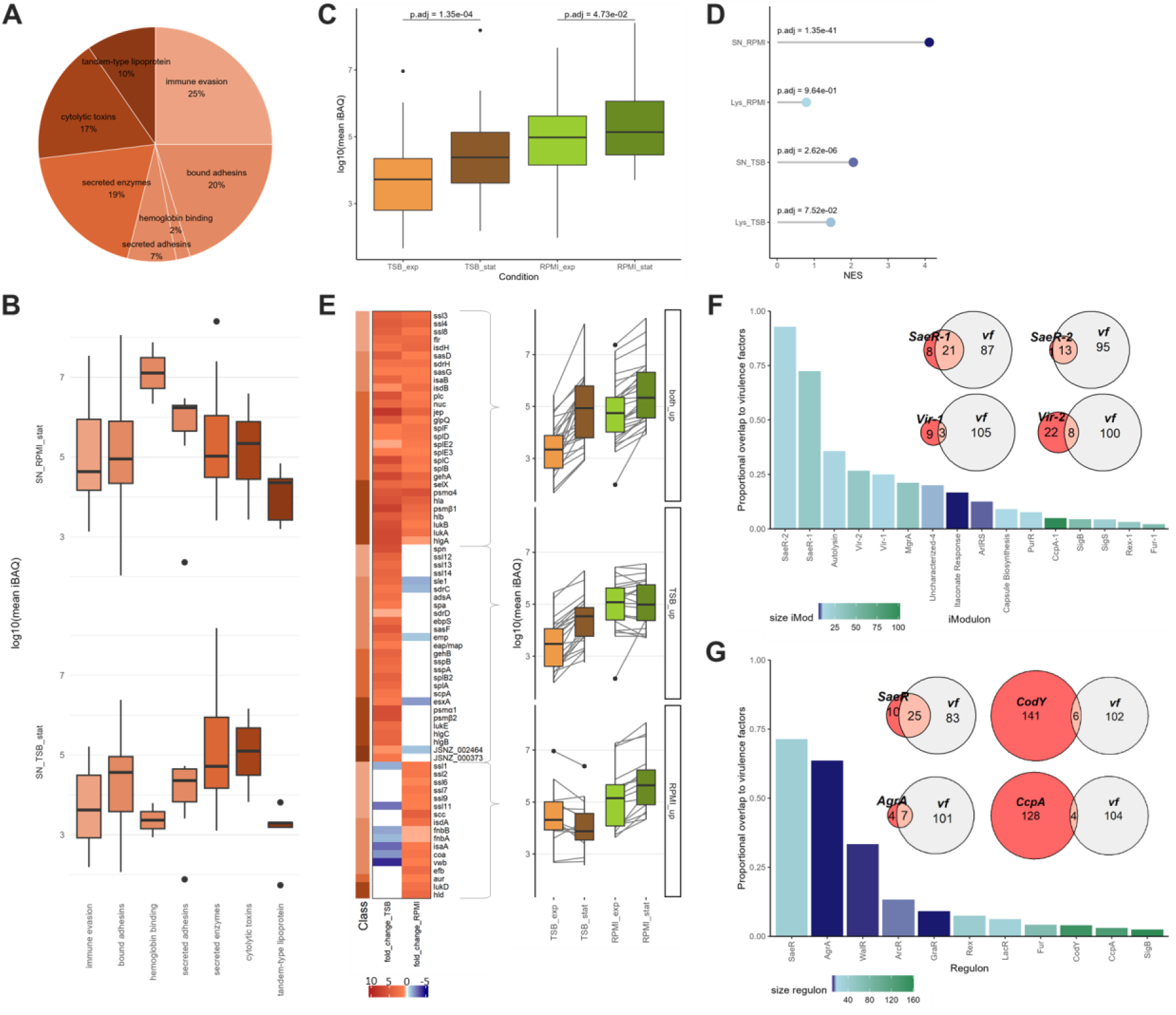
JSNZ virulence factor analysis. **(A)** Composition of virulence factors encoded in the JSNZ genome, classified according to Table S2. **(B)** Comparative abundances of virulence factor classes in supernatants of stationary phase cultures grown in TSB and RPMI. Class colour coding corresponds to panel (A). **(C)** Increase in virulence factor abundance in the culture supernatant from exponential to stationary growth phase, grown in TSB or RPMI. Protein abundances are depicted by mean OD-normalised iBAQ. Benjamini-Hochberg adjusted p-values are derived from pairwise t-test. **(D)** Enrichment of virulence factors in the stationary growth phase compared to exponential growth. Normalised enrichment scores (NES) derived from GSEA statistics, point colour indicates adjusted p-values (Benjamini-Hochberg). **(E)** Heatmap of fold-changes in virulence factor abundance from exponential to stationary growth phase, filtered for significant increase (absolute fold-change >1.5; adjusted p-value <0.05) and reliable detection in at least one growth phase (iBAQ ranked quantile >3). Grouping is based on observed pattern; group-wise abundances are shown as box plots on the left. Class annotations are indicated by the colour bar on the left, consistent with panel (A). **(F)** Overlap of virulence factors (vf) with iModulons and **(G)** regulons. Bar charts display proportional overlaps relative to iModulon/regulon size (set size >5); bar colour reflects set size. Exemplary intersections are illustrated by Venn diagrams. All quantitative values and statistical analyses are based on four bioreplicates.

Overall, there was a significant increase in the abundance of virulence factors in the exoproteome of stationary phase cultures compared to exponential growth, regardless of the medium (pairwise t-test; Fig. 5C & D). In accordance with a more pronounced increase in TSB medium, the majority of virulence factors was significantly increased in TSB and RPMI (29) or TSB only (27). Nevertheless, a set of 17 virulence factors was significantly increased in RPMI only, including the adhesins FnbA, FnbB, and IsaA, and the clotting factors Coa and vWbp, which were even decreased in TSB (Fig. 5E).

When matching the virulence factors to regulons and iModulons, an expected and concordant notable overlap was confirmed to regulons IcaR, SaeR, AgrA and WalR on the one hand, and iModulons SaeR-1/-2, AgrA, Autolysin, Vir-1/-2 and MgrA on the other hand (Fig. 5F & G). Especially the SaeR sets (iModulons and regulons) exhibited an enrichment pattern in the supernatant that was consistent with the aforementioned virulence factor abundances (significant enrichment during stationary growth phase). In TSB this was accompanied by a significant enrichment of iModulon Vir-1 (e.g. Nuc, ScpA, SpA, UreBCDEG; Fig. 4A). Furthermore, iModulons SaeR-1/-2, Autolysin and MgrA as well as the regulon WalR were more enriched in TSB then in RPMI (Fig. 4B). Interestingly, a different pattern emerged in the cellular proteome: iModulon Vir-2 (e.g. Aur, Ebh, T7SS components) was enriched during the exponential growth phase in RPMI compared to stationary growth in RPMI (Fig. 4A), as well as compared to exponential growth in TSB (Fig. 4B). Similarly, the SaeR regulon and the SaeR-1 iModulon were intracellularly enriched in RPMI compared to TSB during exponential growth.

It is noteworthy that AgrA, one of the best-known virulence regulators in *S. aureus* (Singh and Ray, 2014), although encoded in the genome did not show up in any of the analyses.

### 3.8 Jep and the related Spls

Across all analyses, the protease Jep was found to be a salient feature of JSNZ. Of particular note is the exceptionally high amount of secreted Jep during stationary growth phase (Fig. 6A). Accordingly, we hypothesise a central role for Jep in JSNZ’s physiology and its interaction with the host, thus prompting a particular interest in Jep.

**Figure 6.**
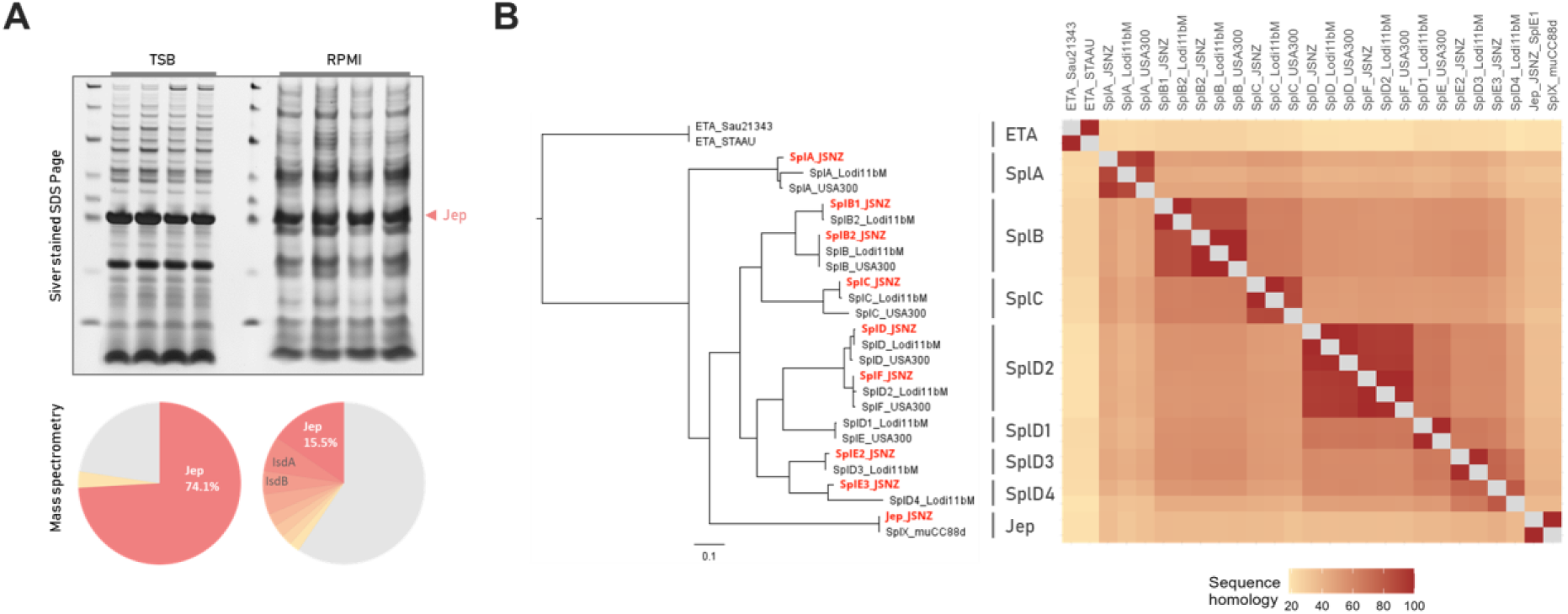
Jep is a highly abundant Spl-like serine protease secreted by JSNZ. **(A)** Composition of stationary-phase JSNZ culture supernatants grown in TSB (left) or RPMI (right) shown *via* silver-stained SDS-PAGE (top) and iBAQ-based mass spectrometry quantification (bottom; four bioreplicates). Jep is by far the most abundant protein in both media. **(B)** Protein sequence distance tree (left) and similarity matrix (right) comparing Jep and the eight Spls encoded by JSNZ (highlighted in red) with the Spls from *S. aureus* strains USA300_FPR3757 and Lodi11bM, as well as SplX from muCC88d and two ETA sequences. USA300 (CC8) was selected as established reference strain; Lodi11bM (CC126) for its comparable νSaβ island structure to JSNZ. Spl nomenclature follows Kläui *et al*. (2019).

Jep is a secreted serine protease, that is encoded within a Sa1int prophage and in this respect analogous to Eta (Fig. 1C). However, the amino acid sequence similarity and phylogeny revealed a close relationship to the Spls (Fig. 6B). They shared a conserved catalytic triad and a comparable signal peptide for Sec-dependent secretion (Fig. S10). The high sequence homology furthermore indicates an overall common protein structure with the Spls, Eta and also the V8 protease SspA (Dubin *et al*., 2008).

In addition to Jep, the JSNZ genome encodes eight Spl variants (Fig. S6). These include variants clearly assigned as SplA, SplB (two copies), SplC, SplD, and SplF, based on amino acid sequence similarities exceeding 85% compared to the corresponding variants in USA300 (Fig. 6B). The remaining two Spls were annotated as SplE variants (SplE2, SplE3), although they exhibited a greater divergence from the USA300 SplE (65% and 64%, respectively). Given that the JSNZ νSaβ island resembles the νSaβ type XV, the JSNZ Spl variants were further compared to the nine Spl variants encoded by the respective type XV reference strain Lodi11bM (Kläui *et al*., 2019). Notably, SplE2 and SplE3 from JSNZ clustered with SplD3 and SplD4 from Lodi11bM, respectively. In light of these findings, a re-annotation following the proposed nomenclature of Kläui *et al*. (2019; SplD -> SplD2, SplF -> SplD2, SplE2 -> SplD3, SplE3 -> SplD4) may be warranted.

While present in an impressive amount, Jep replicates the same regulation pattern as the Spls under standard laboratory conditions, i.e. a strong increase in abundance of the proteases in the supernatant towards the stationary phase (Fig. 5E). However, the genetic localisation is fundamentally different and no regulatory sequence was mapped upstream of *jep* in our genome-wide analysis.

## 4 Discussion

### 4.1 JSNZ is a good representative for mouse-adapted *S. aureus* isolates

JSNZ meets many of the hallmarks of a reference strain for murine *S. aureus* infection models. JSNZ belongs to CC88, which is rarely identified in human isolates (Holtfreter *et al*., 2016) but is by far the predominant lineage among *S. aureus* isolates from laboratory mice (∼50%; Mrochen *et al*., 2018a; Yebra *et al*., 2024). Additionally, the reconstructed phylogeny exhibits a low genetic diversity within the international murine CC88 lineage, which likely originated from a single human-to-mouse transmission followed by global spread through commercial vendors (Yebra *et al*., 2024). This makes CC88 a compelling model to investigate the adaptation from the human to the murine host. However, while some murine CC88 isolate still show genetic features common in human isolates but rare in murine isolates (Schulz *et al*., 2017), JSNZ carries none of these features. It lacks most superantigens except for *selX* and *sel26*, antibiotic resistances (e.g. *blaZ* and *mecA*), and prophage Sa3int with the known immune evasion cluster 1 (IEC1; *scn*, *chp*, *sak*). The combination of familiar host-specific features, a suitable infection phenotype in mice, and relative ease of genetic manipulation (Chan *et al*., 2024; Holtfreter *et al*., 2013) explains why JSNZ is used in various infection studies.

Nevertheless, despite the initial characterisation of genetic traits (Holtfreter *et al*., 2013), a complete, closed genome sequence and a resulting proteome profile remain missing. Therefore, the JSNZ reference genome and baseline proteome profiles provided in this study strengthen the knowledge base on this increasingly important model strain and advance its characterisation as a reference strain for mouse infection models. The implementation in *Aureo*Wiki ensures broad accessibility and cross-linking with genome information of other *S. aureus* reference strains.

### 4.2 Deletion in HsdR facilitates genetic modification

The described efficient transformability of the CC88 JSNZ with plasmids derived from the CC8 strain RN4220 (Holtfreter *et al*., 2013) is likely based on the deletion in *hsdR* uncovered in this study. HsdR is the restriction subunit of the Sau1 type I restriction-modification (Sau1RM) system, which is the basis for the lineage-specific transformation barrier present in *S. aureus* (Roberts *et al*., 2013; Waldron and Lindsay, 2006). Its truncation in RN4220 (W197*) broke down the barrier, establishing this strain as the most important host for genetic engineering in *S. aureus* (Nair *et al*., 2011). Although, the deletion in JSNZ’s HsdR (Δ705-876) is substantially smaller than the inactivating truncation in RN4220, it disrupts the CTD. The CTD of HsdR, in turn, is essential for the assembly of the HsdR_2_M_2_S_1_ holoenzyme, and deletion of this domain abolishes the DNA-cleavage activity of the system *in vitro* (Gao *et al*., 2020). Nevertheless, since the two pairs of HsdM and HsdS are intact, an active methyltransferase M_2_S_1_ can be formed and introduce CC88 specific DNA-methylation. Consequently, JSNZ is the gateway to the genetic modification of CC88 strains, as is already exemplified by transformation of WU1, which cannot be directly transformed with RN4220-derived plasmids (Bobrovskyy *et al*., 2023). Of note, in contrast to RN4220, JSNZ retains an active SauUSI restriction enzyme (JSNZ_002468). SauUSI blocks the direct introduction of cytosine-methylated DNA, e.g. yielded from common *dcm*^+^ *E. coli* strains (Corvaglia *et al*., 2010; Loenen and Raleigh, 2014; Xu *et al*., 2011).

### 4.3 Genetic traits indicate adaptation to the murine host

The virulence gene profile of JSNZ matches well with the colonising phenotype of this strain with comparably low virulence for laboratory mice. We identified a total of 108 virulence genes in the JSNZ genome, 99 of which were expressed under laboratory conditions. Their functional distribution was dominated by immune evasion, adhesion and clotting (Fig. 5A). A similar composition is associated with *S. aureus* strains causing low cytotoxicity but high within-herd prevalence in dairy cows (Addis *et al*., 2022). The observed virulence factor composition and secretion may thus favour the wide dissemination of JSNZ and related CC88 strains in laboratory mice (Sun *et al*., 2018; Trübe *et al*., 2019). In addition, a number of virulence factors showed up with potential lineage-specific amino acid sequence difference between CC8 and CC88, including factors of immune evasion, adhesion and agglutination (Fig. 2A). These factors may be promising candidates to evaluate the proposed lineage-specific pre-adaptation of CC88 to the murine host (Yebra *et al*., 2024). For example, the CC88 variants of Coa and vWbp lead to reduced agglutination of human plasma, but at least slightly increased agglutination of mouse plasma compared to the CC8 reference variants (Sun *et al*., 2018; Yebra *et al*., 2024).

A noticeable characteristic on a genomic scale is the Sa1int prophage. The φLabRodCC88_3 carried by JSNZ was just recently described as the most frequent prophage in murine isolates, harbouring several motifs enriched in the genomes of rodent *S. aureus* isolates compared to human isolates (Yebra *et al*., 2024). This is also reflected in our data in the accumulation of unique genes in the region of φLabRodCC88_3 compared with the *Aureo*Wiki genomes (Supplementary data 1: Table 1-1, Supplementary data 4). Among these unique genes, the serine protease Jep is of particular interest due to its exceptionally high abundance in the exoproteome, underscoring its potential relevance for JSNZ. Considering the close relatedness of Jep to the Spls we suspect a comparable function. There is evidence that the expression of Spls is upregulated under skin-like conditions (Costa *et al*., 2024). Upon invasion of the host, they may promote spreading (Paharik *et al*., 2016), contribute to immune evasion (Dasari *et al*., 2022; Scherr *et al*., 2024) and trigger allergic responses (Stentzel *et al*., 2017; Teufelberger *et al*., 2018).

The overall diversity of Spls, including Jep, may be a genetic trait that facilitates host switches. The νSaβ type most similar to that of JSNZ – type XV with nine encoded *spls* – was first identified in bovine *S. aureus* CC126 isolates (Kläui *et al*., 2019). Just like CC88, CC126 is an example for a successful host jump with subsequent expansion of the pathogen (Yebra *et al*., 2022). Since each Spl variant has a specific target sequence (Stach *et al*., 2021), it is plausible that the other Spl variants and Jep might possess new host-related specificities. Elucidating the function of Spls and Jep could therefore also contribute to our understanding of lineage-dependent host adaptation (Matuszewska *et al*., 2020).

The absence of a Sa3int prophage has already been described for a number of *S. aureus* isolates from rodents (Mrochen *et al*., 2018a, 2018b; Raafat *et al*., 2020; Sun *et al*., 2018) and also from other mammalian hosts (Hau *et al*., 2015; Price *et al*., 2012; Resch *et al*., 2013; Spoor *et al*., 2013). A possible role of the resulting active Hlb for long-term establishment in animal hosts has been extensively reviewed (e.g., Rohmer and Wolz, 2021). Among other aspects, active Hlb in mice contributes to *S. aureus* colonisation (Jia *et al*., 2023; Katayama *et al*., 2013) and infection-associated hypoxia (Li *et al*., 2025). Interestingly, in addition to active Hlb, the two analysed CC88 mouse isolates exhibited a mutated Hla which is expressed but not haemolytically active (Trübe *et al*., 2019). In mouse models, Hla inactivation leads to reduced severity of infection of the skin (Kennedy *et al*., 2010), eyes (Putra *et al*., 2019), brain (Kielian *et al*., 2001), breast (Jonsson *et al*., 1985), and lungs (Tang *et al*., 2019). However, in an intraperitoneal model, it has been shown that although lethality is reduced, abscess formation is unchanged (Rauch *et al*., 2012). Interestingly, also the regulator Rbf is truncated in both murine CC88 strains. Although Rbf is mainly known as a regulator of biofilm formation *via* IcaR (Cue *et al*., 2013), it is also a potent repressor of *hla* and *psmα* (Fang *et al*., 2020). This would be in line with the absence of IcaABCD in our proteomic analyses on the one hand, and the relatively high abundance observed for Hla on the other hand (Fig. S7). Hence, the mutations in *hla* and *rbf* might be correlated. Taken together, it is possible that the active Hlb to some degree balances the Hla inactivation to ensure host-adapted virulence that favours long-term colonisation and thus transmission of *S. aureus*. Amino acid sequence differences of many surface proteins including Ebh are known to be lineage-specific (McCarthy and Lindsay, 2010) and their variation is often part of the adaptation to new hosts (Herron-Olson *et al*., 2007; Monecke *et al*., 2025). In fact, the relatively large variability of Ebh was suggested as a mechanism for evolutionary adaptation introduced by the nearby integration site of Sa2int phages (Chen *et al*., 2018).

### 4.4 Medium and growth phase strongly influence the proteome

Besides the completed genome, we also provide a comprehensive proteomic data set for JSNZ cultivated under standard laboratory conditions. By including exponential and stationary growth phase as well as the cellular proteome and exoproteome, this data set is a valuable resource to assess the expected physiological state and expressed proteome composition when working with JSNZ under defined experimental conditions. This is important, as expression pattern and proteome profile are known to change massively between different growth states (Becher *et al*., 2009; Liebeke *et al*., 2011; Mäder *et al*., 2016).

Overall, the proteome of JSNZ in complex and synthetic medium is comparable to that of other *S. aureus* strains. The proteins that are consistently highly abundant in our data set correspond to the highly expressed genes in *S. aureus* strain NCTC8325 (Mäder *et al*., 2016) and widely overlap with essential genes determined in the NCTC 8325-derivate SH1000 (Chaudhuri *et al*., 2009). In TSB, JSNZ furthermore clearly revealed the change from a growing state with high level of bacterial replication and protein synthesis to a non-growing state with a diversified energy metabolism and elevated stress responses (Becher *et al*., 2009; Kohler *et al*., 2005). This is mainly due to the exhaustion of the primary carbon source glucose and further nutrients (Liebeke *et al*., 2011; Poudel *et al*., 2020).

Cultivation in RPMI fundamentally changes the situation (Liebeke *et al*., 2011). Significant enrichment of Fur-dependent proteins during cultivation in RPMI compared to TSB accounts for the lack of available iron in RPMI (Xiong *et al*., 2000). Additionally, the PurR regulon is significantly enriched in RPMI compared to TSB during exponential growth phase (Fig. 4B). This is explained by the lack of purines in RPMI medium, resulting in an increase of intracellular phosphoribosyl pyrophosphate (PRPP) and downstream derepression of PurR regulated genes (Grove, 2025). The stationary phase is characterised by a severe shortage of most nutrients and induction of the stringent response (Carrilero *et al*., 2023; Poudel *et al*., 2020). Under these conditions, (p)ppGpp outcompetes PRPP in binding to PurR and repression is restored (Anderson *et al*., 2022; Salzer *et al*., 2025). In TSB, purines are present and scavenged from the medium. Consequently, PurR dependent proteins only increase in abundance when purines are fully depleted with onset of the stationary phase (Fig. 4A).

### 4.5 Influence of medium choice on virulence factor expression

In both media, we observed a significant increase in the level of virulence factors towards stationary growth phase. This applies to the absolute abundances as well as the relative proportion in the stationary phase exoproteome (Fig. 5C&D). Nevertheless, we observed a different pattern of virulence factor expression in TSB and RPMI, respectively, in line with observations of increased adhesion of *S. aureus* grown in RPMI (Wijesinghe *et al*., 2018).

Regulation of virulence factors in most cases is a combination of responses to a plethora of regulatory signals (King *et al*., 2020; Le Scornet and Redder, 2019; Poudel *et al*., 2020). In general, our data suggests a strong influence of SaeR on virulence gene expression also in JSNZ (Fig. 4A). SaeR is one of the major virulence regulators in *S. aureus*, influencing virulence factors of different functions (Geiger *et al*., 2008; Liu *et al*., 2016; Mainiero *et al*., 2010). Accordingly, there are two differing iModulons assigned to SaeR: SaeR-1 mainly contains adhesins and immune evasion factors, while SaeR-2 contains the secreted proteases, leukocidins and staphylococcal protein A (*S. aureus* Precise165 Dataset, Poudel *et al*., 2022). An important part of these differentiating adjustment is the influence of *S. aureus’* metabolic state on the virulence expression. To highlight just a few interconnections: CodY also represses *sae* expression (Mlynek *et al*., 2018), PurR also represses *sarA*, *fnbA* and *ssl11* expression (Goncheva *et al*., 2020; Sause *et al*., 2019), and CcpA increases RNAIII levels as well as contributes to transcription regulation of e.g. *spa* and *hla* (Seidl *et al*., 2006). These regulatory cross-links, for example, aggravate *S. aureus* infections in glucose-rich niches such as the liver or in diabetic mice (Bischoff *et al*., 2017). Furthermore, PurR was recently shown to interact with T7SS targets (Bronesky *et al*., 2019).

The dominance of toxins and secreted enzymes in the stationary exoproteome of JSNZ grown in TSB is thus likely linked to the depletion of glucose, amino acids and nucleotides from the medium. In RPMI, the inactivation of PurR and CodY during exponential phase explains the early peak of Vir-1 (T7SS) and SaeR-1 (adhesins), that, however, is only detected intracellularly (Fig. 4A). The fact that the SaeR regulon overall is nevertheless significantly enriched during stationary phase in RPMI is most likely due to the induction of virulence gene expression by iron deficiency (Busch *et al*., 2025; Johnson *et al*., 2011; Torres *et al*., 2010).

The virulence pattern observed in RPMI is of particular interest, since RPMI is frequently used in the preparation of infection experiments and to approximate host environments *in vitro* (Dörries and Lalk, 2013; Hofstee *et al*., 2020; Li *et al*., 2024; Miajlovic *et al*., 2010; Zapotoczna *et al*., 2015). In fact, *S. aureus* grown in human serum also exhibits increased expression of CodY dependent genes, transient upregulation of PurR dependent genes and a strong increase of Fur regulated genes (Mäder *et al*., 2016; Malachowa *et al*., 2011; Poudel *et al*., 2020). Heme and iron uptake is essential for the survival of *S. aureus* in the bloodstream (Bleackley *et al*., 2009; Skaar and Schneewind, 2004). However, the also described downregulation of Agr dependent genes (Poudel *et al*., 2020) is not observed in in JSNZ grown in RPMI. The Agr activity is not altered significantly in our data. The same applies to skin infections, which are additionally characterised by a downregulation of SigB-dependent genes (Poudel *et al*., 2020) as well as an upregulation of the *fadXDEBA* and the urease operon (Enroth *et al*., 2025). None of these effects could be observed in the proteome of JSNZ grown in RPMI.

These comparisons once again emphasise the great importance of nutrient availability as a signal for the infection status (Balasubramanian *et al*., 2017). In this respect, the composition of RPMI also in JSNZ induces a proteome pattern that is roughly comparable to that expected from growth in serum. Nevertheless, it becomes apparent that the regulation of virulence factors is additionally dependent on specific host signals missing in RPMI. Accordingly, the virulence factor equipment shows specific differences between RPMI and host environment that must be considered when interpreting experimental outcomes.

## 5. Conclusion

In this study, we provide a comprehensive genomic and proteomic data set to advance the characterisation of JSNZ as reference strain for mouse-adapted *S. aureus*. This includes publication of the reannotated genome sequence of JSNZ on *Aureo*Wiki, which allows a user-friendly comparison with established *S. aureus* strains. Based on our data, we could explain the good transformability of JSNZ due to a deletion in HsdR. In addition, the molecular basis for the colonising phenotype of JSNZ and possible adaptations to the murine host become apparent. Of note, in the restricted comparison with selected other CC88 strains, this lineage appears to be a promising candidate for further investigation of host adaptation. While the proteomic profile of JSNZ proves its comparability to established *S. aureus* reference strains, an exceptionally high amount of the secreted serine protease Jep is particularly striking. Consequently, the role of Jep in bacterial physiology and during infection in murine hosts is of particular interest and may allow conclusions to be drawn about the function of the closely related Spls.

## Acknowledgements

We would like to thank Alexander Ganske, Anja Wiechert, and Katrin Schoknecht for their excellent assistance with some experiments, and Katrin Stark for her administrative support.

## CRediT authorship contribution statement

**Hannes Wolfgramm**: Conceptualization, Data curation, Formal analysis, Investigation, Methodology, Visualization, Writing – original draft, Writing – review & editing. **Larissa M. Busch**: Data curation, Formal analysis, Investigation, Methodology, Writing – original draft, Writing – review & editing. **Jöran Tebben**: Investigation, Writing – review & editing. **Henry Mehlan**: Data curation, Formal analysis, Methodology, Software, Writing – review & editing. **Lisa Hagenau**: Methodology, Writing – review & editing. **Thomas Sura**: Methodology, Writing – review & editing. **Tilly Hoffmüller**: Investigation, Writing – review & editing. **Elisa Bludau**: Investigation, Writing – review & editing. **Manuela Gesell Salazar**: Investigation, Data curation, Writing – review & editing. **Alexander Reder**: Methodology, Writing – review & editing. **Stephan Michalik**: Methodology, Software, Writing – review & editing. **Leif Steil:** Methodology, Software, Writing – review & editing. **Kristin Surmann**: Funding acquisition, Supervision, Writing – review & editing. **Ulrike Mäder**: Data curation, Resources, Writing – original draft, Writing – review & editing. **Silva Holtfreter**: Conceptualization, Funding acquisition, Supervision, Writing – original draft, Writing – review & editing. **Uwe Völker**: Conceptualization, Funding acquisition, Project administration, Supervision, Writing – review & editing.

## Declaration of generative AI and AI-assisted technologies in the writing process

During the preparation of this work the authors used Microsoft Copilot and DeepL in order to improve the readability and language of the manuscript. After using these tools, the authors reviewed and edited the content and take full responsibility for the content of the published article.

## Data availability

The annotated genome has been submitted to NCBI and is available for exploration *via Aureo*Wiki (aureowiki.med.uni-greifswald.de). The spectral libraries and MS analysis output data have been deposited to the ProteomeXchange Consortium *via* the PRIDE (Perez-Riverol *et al*., 2025) partner repository with the dataset identifier PXD067993.

