## Supplementary figures and images for "Integrated genomic and proteomic analysis of the mouse-adapted *Staphylococcus aureus* strain JSNZ"

### Supplementary Data 4

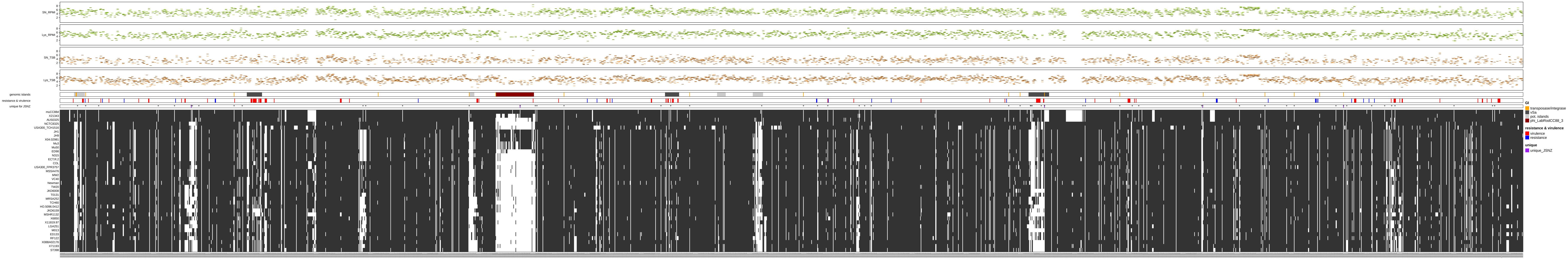
