## Supplementary Figures and Tables for "Integrated genomic and proteomic analysis of the mouse-adapted *Staphylococcus aureus* strain JSNZ"

**This PDF includes:**

**Figure S1.** Genomic comparison of genomic islands (GI), island-like structures and repetitive gene clusters in the JSNZ genome to the four other CC88 genomes considered in this study.

**Figure S2.** Protein sequence alignment of cell-wall-anchored protein SasG orthologs.

**Figure S3.** Assignment of the two JSNZ encoded HsdS (JSNZ_000367 and JSNZ_001801) to the respective target recognition domains (TRDs).

**Figure S4.** Schematic visualization of the novel Type IV restriction system, classified as a coiled-coil tandem (CoCoNuT) of subtype I-B.

**Figure S5.** Abundances of proteins identified in JSNZ exponential and stationary cultures grown in TSB or RPMI.

**Figure S6.** Proteomic coverage of genomic islands and island-like structure depicted by protein abundance assigned to corresponding genes.

**Figure S7.** Abundance of detected proteins encoded by annotated pseudogenes in the JSNZ genome.

**Figure S8.** Enrichment of protein sets comparing growth phases.

**Figure S9.** Enrichment of protein sets comparing media.

**Figure S10.** Protein sequence alignment of Jep to the Spls encoded in JSNZ and USA300_FPR3757.

**Table S1.** Summary of sequencing read statistics.

**Table S2.** Reference integrase sequences used for phage classification.

**TableS3.** Information on reversed phase liquid chromatography (RPLC).

**TableS4.** Information on data independent analysis (DIA) mass spectrometry.

**Additional files:**

**Supplementary data 1.** Information on annotated genes, corresponding orthologs, detected abundance of corresponding proteins and statistics.

- **Supplementary table 1-1.** Curated annotation of the JSNZ genome including meta information.
- **Supplementary table 1-2.** Regulon prediction of annotated JSNZ genes.
- **Supplementary table 1-3.** Protein sequence similarity of the annotated JSNZ genes to orthologous genes of *S. aureus* strains muCC88d, AUS0325, 21343, COL, NCTC8325, Newman, and USA300.
- **Supplementary table 1-4.** Mass spectrometric detected and quantified proteins in JSNZ culture samples grown in TSB and RPMI.
- **Supplementary table 1-5.** Consistently high-abundant proteins of JSNZ.
- **Supplementary table 1-6.** ROPECA statistics on protein alterations between exponential and stationary growth phase per medium.
- **Supplementary table 1-7.** ROPECA statistics on relative protein alterations between JSNZ grown in TSB and RPMI per growth phase.

**Supplementary data 2.**

- **Supplementary table 2.** Overview on JSNZ genes classified as virulence or resistance related.

**Supplementary data 3.** Information on occurance of mouse-specific mutations in JSNZ, peptide coverage of detected proteins assigned to pseudogenes and proteins with unexpected localization.

- **Supplementary table 3-1.** Occurrence of mouse-associated, non-phage located mutations in the JSNZ genome.
- **Supplementary table 3-2.** Proteins with unexpected distribution between cellular fraction and supernatant contradicting predicted localization (DeepLocPro).
- **Supplementary table 3-3.** Peptide coverage of the five pseudogene derived proteins detected by mass spectrumetry.

**Supplementary data 4.** Graphical summary of the JSNZ genome, orthologous genes in the AureoWiki strains, and proteomic coverage across all conditions.

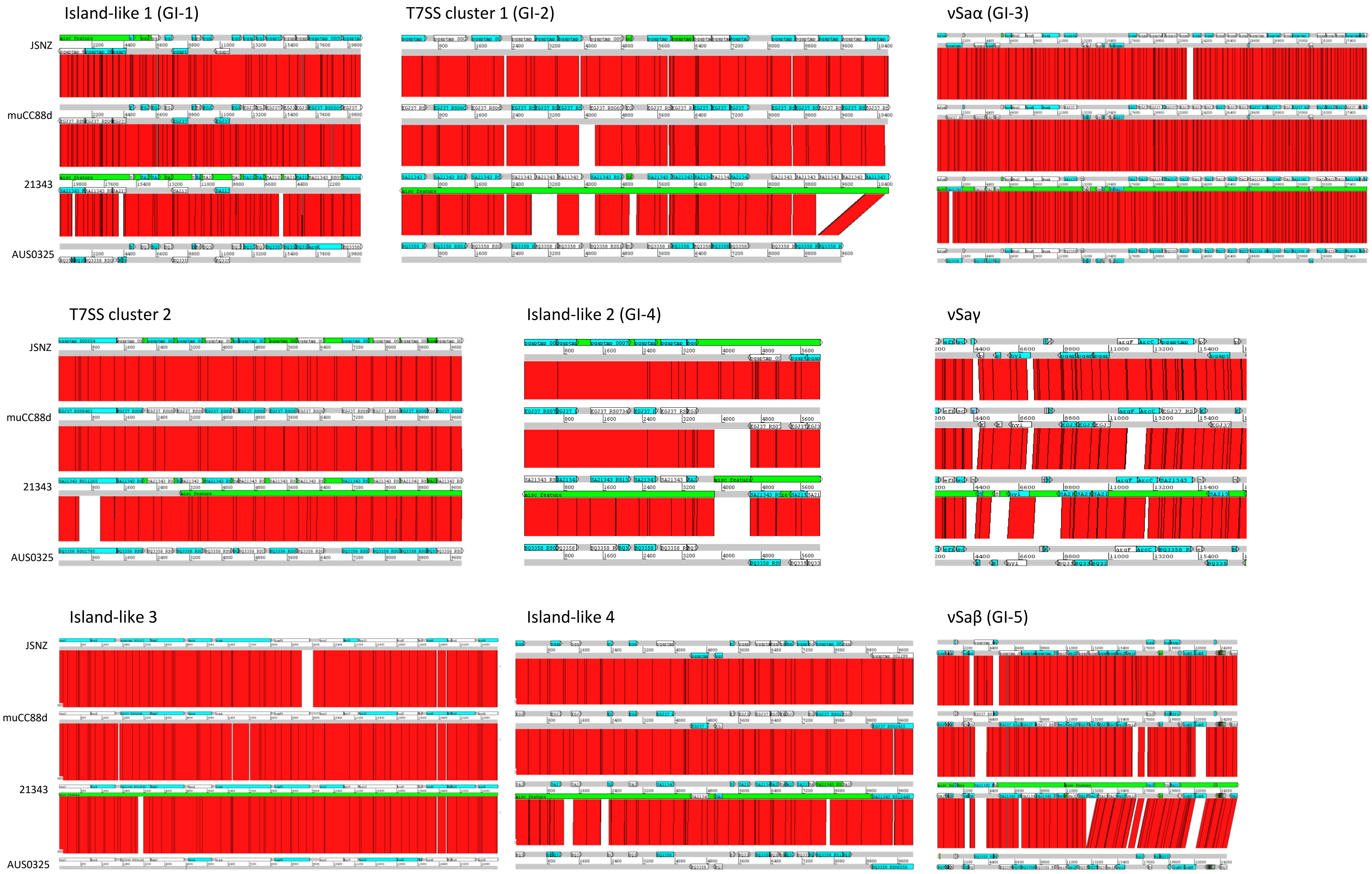


**Figure S1.** Genomic comparison of genomic islands (GI), island-like structures and repetitive gene clusters in the JSNZ genome to the three other CC88 genomes considered in this study. Genomic structure was compared with the Artemis Comparison Tool, identity level was set to 95%. GIs defined as specific for CC88 according to Kpeli *et al.* (2017) are mentioned in brackets. The comparison of ΦLabRodCC88_3 is depicted in Fig. 1C of the main text.

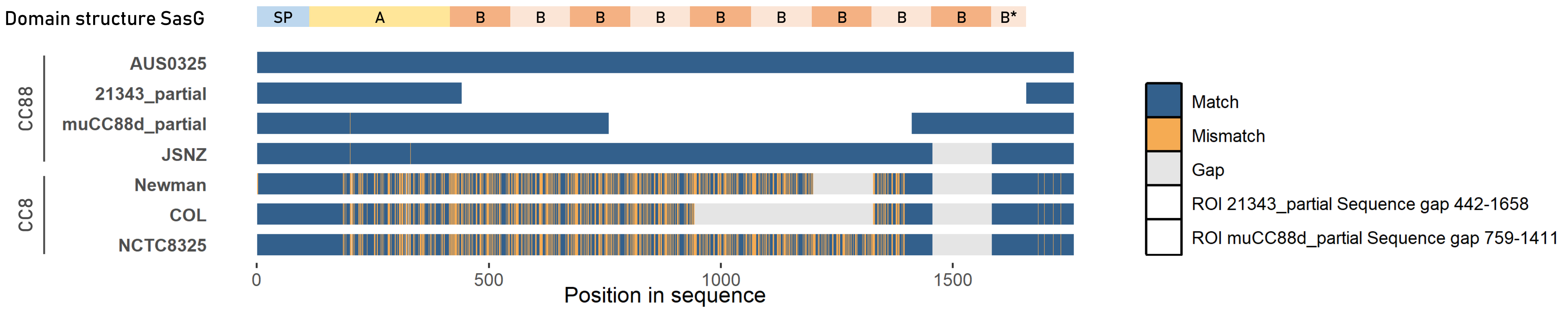


**Figure S2.** Protein sequence alignment of cell wall-anchored protein SasG orthologs. Amino acid mismatches (yellow), gaps (grey) and premature stop codons (dark red) are depicted. Domain structure is indicated above the alignment. Due to the repetitive structure of B domains, short-read genome sequences of isolates 21343 and muCC88d exhibit a sequencing gap interrupting the *sasG* sequence. SP signal peptide, B* truncated B domain.

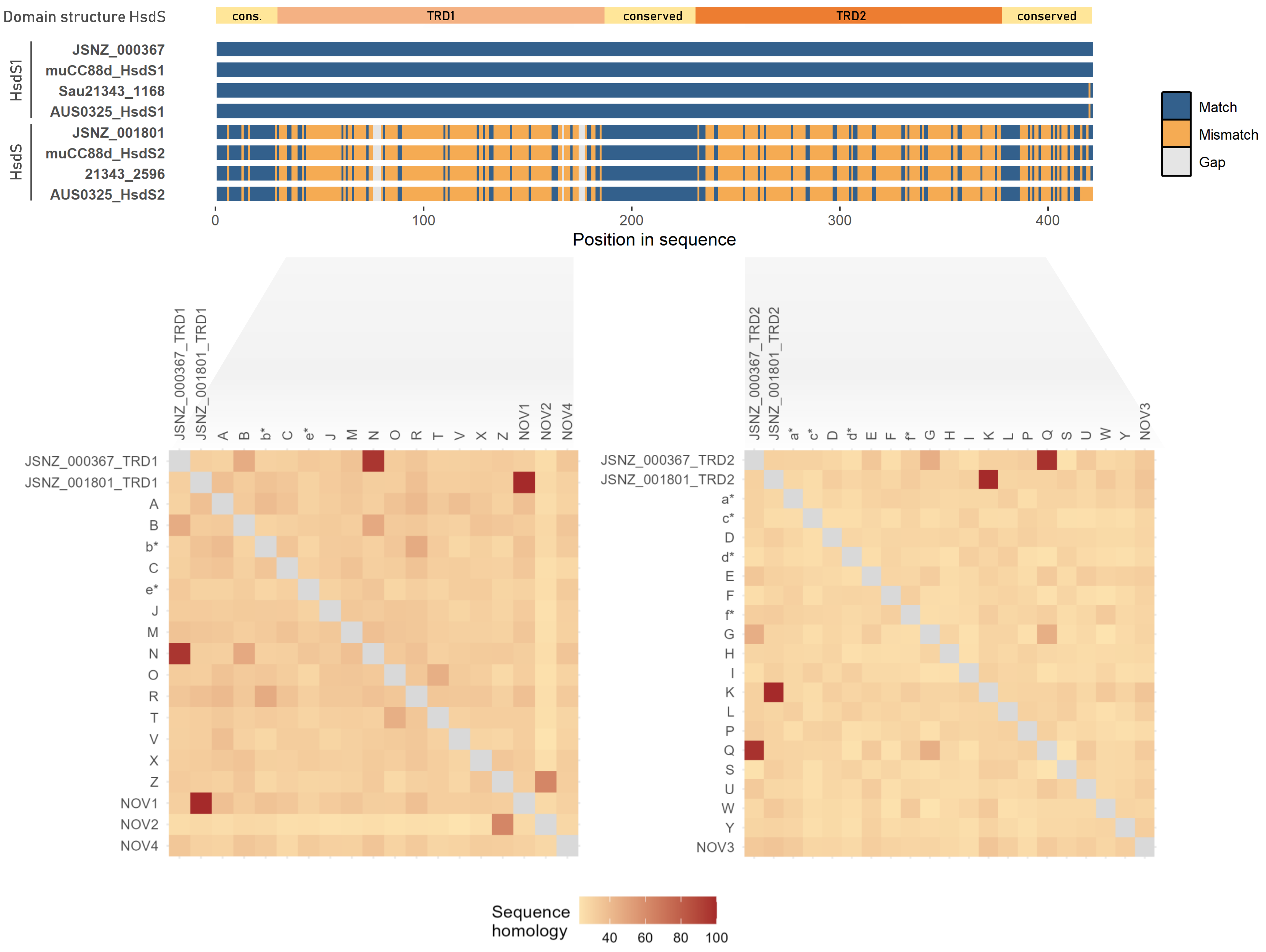


| Locus tag | Annotation | TRDs | Recognition sequence |
|---|---|---|---|
| JSNZ_000367 | HsdS1 | N-Q | ACC-5-RTGT |
| JSNZ_001801 | HsdS | NOVEL1-K | GAG-6-TCG |

**Figure S3.** Assignment of the two JSNZ encoded HsdS (JSNZ_000367 and JSNZ_001801) to the respective target recognition domains (TRDs). (Top) Protein sequence alignment of both HsdS copies and their orthologs in the other analysed CC88 isolates. Protein domain structure is depicted above. (Bottom) Sequence similarity matrix comparing TRD1 and TRD2 of JSNZ_000367 and JSNZ_001801 to the reference sequences published by Cooper *et al.* (2017). Each of the four TRDs can be clearly assigned to a reference TRD, as indicated by the distinct red squares. The resulting TRD combination with according recognition sequence is listed in the table. Underlined nucleotides are methylation targets.

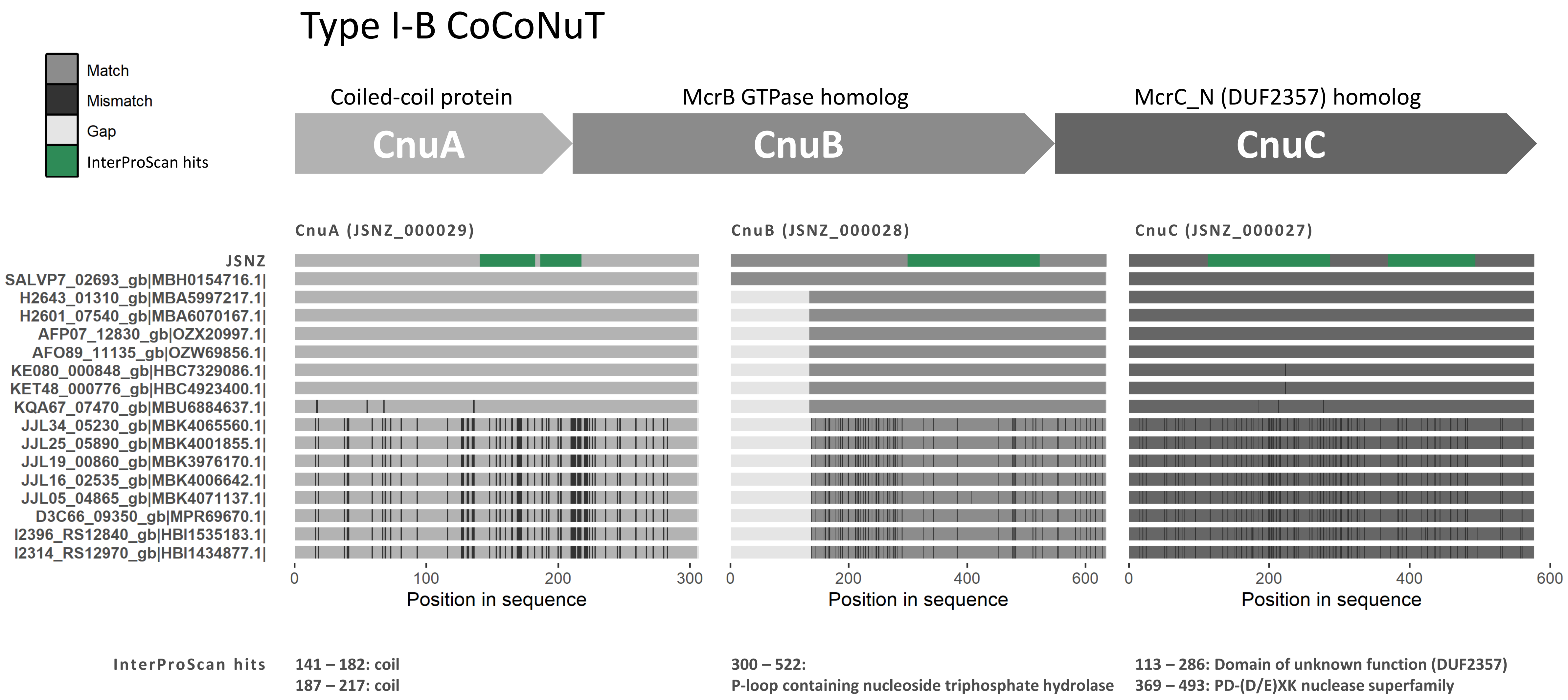


**Figure S4.** Schematic visualization of the novel type IV restriction system, classified as a coiled-coil tandem (CoCoNuT) of subtype I-B. Classification was validated by (i) protein sequence alignment of JSNZ_000027 to JSNZ_000029 with representative CnuA, CnuB and CnuC sequences (Bell *et al.*, 2024) and (ii) identification of typical domains *via* InterProScan analysis, highlighted green in the upper JSNZ sequence and detailed below the alignment.

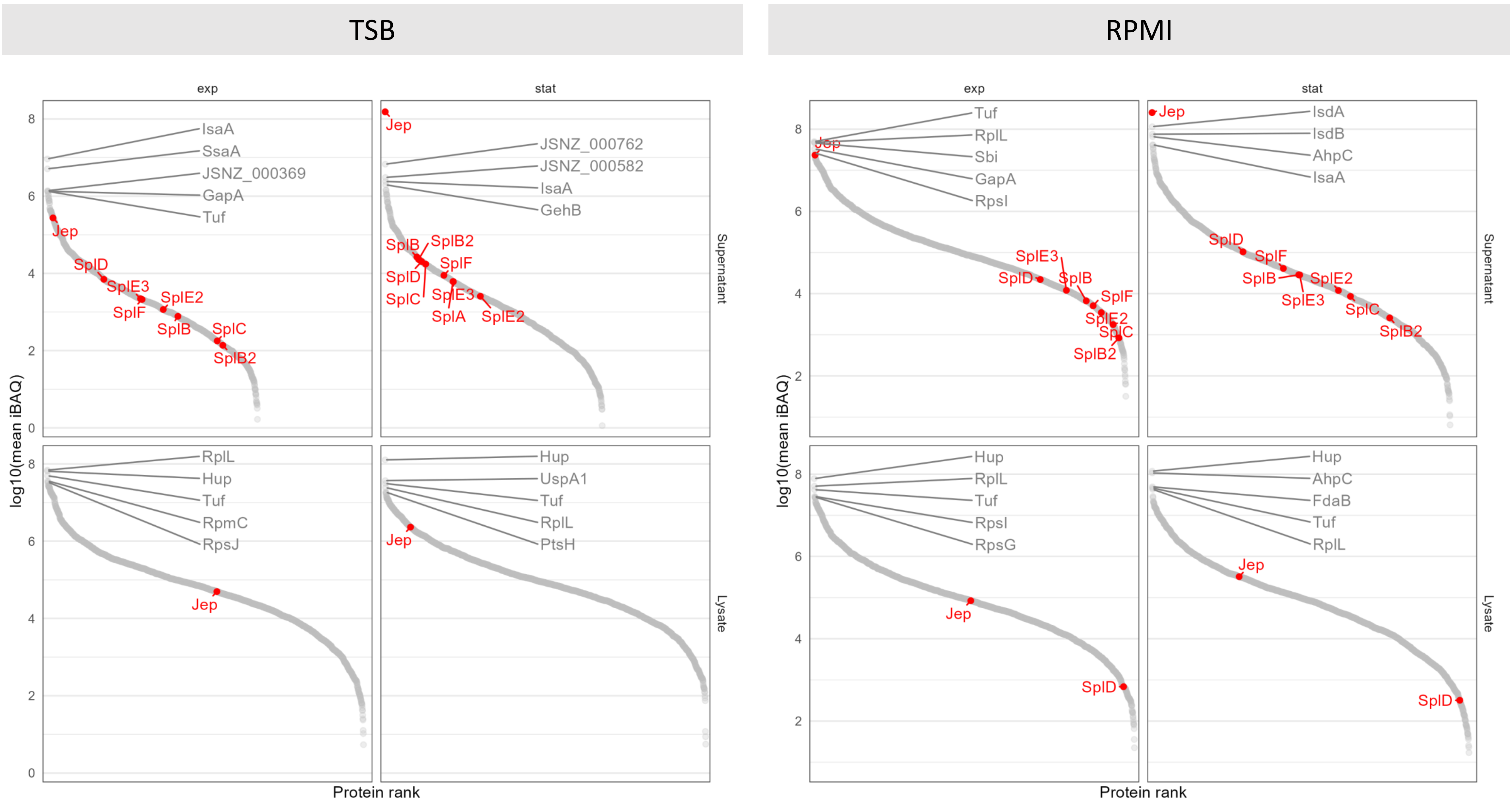


**Figure S5.** Abundances of proteins identified in JSNZ exponential and stationary cultures grown in TSB or RPMI. Proteins separately ranked by iBAQ. Secreted serine-protease like proteins (Spls) and JSNZ extracellular protease (Jep) are labelled in red, top five most abundant proteins per sample are labelled in grey.

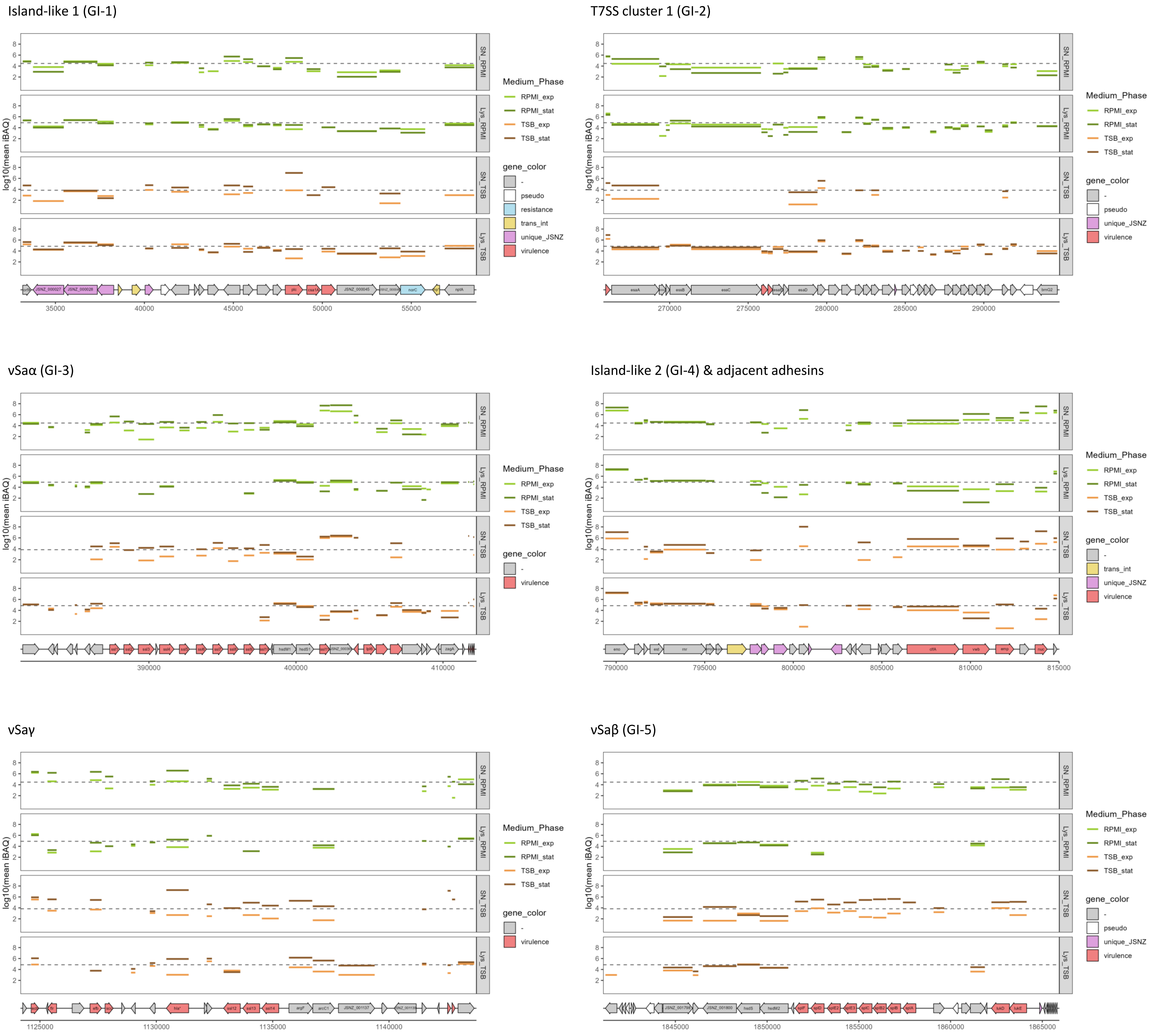


**Figure S6.** Proteomic coverage of genomic islands and island-like structure depicted by protein abundance assigned to corresponding genes. Islands were selected based on contained virulence genes. The annotated genome region is shown with genome position in base pairs. Median iBAQ per medium and fraction is indicated by a dashed line, protein iBAQ lines are coloured based on condition. iBAQ calculation based on four bioreplicates. ΦLabRodCC88_3 is depicted in Fig. 3F of the main text.

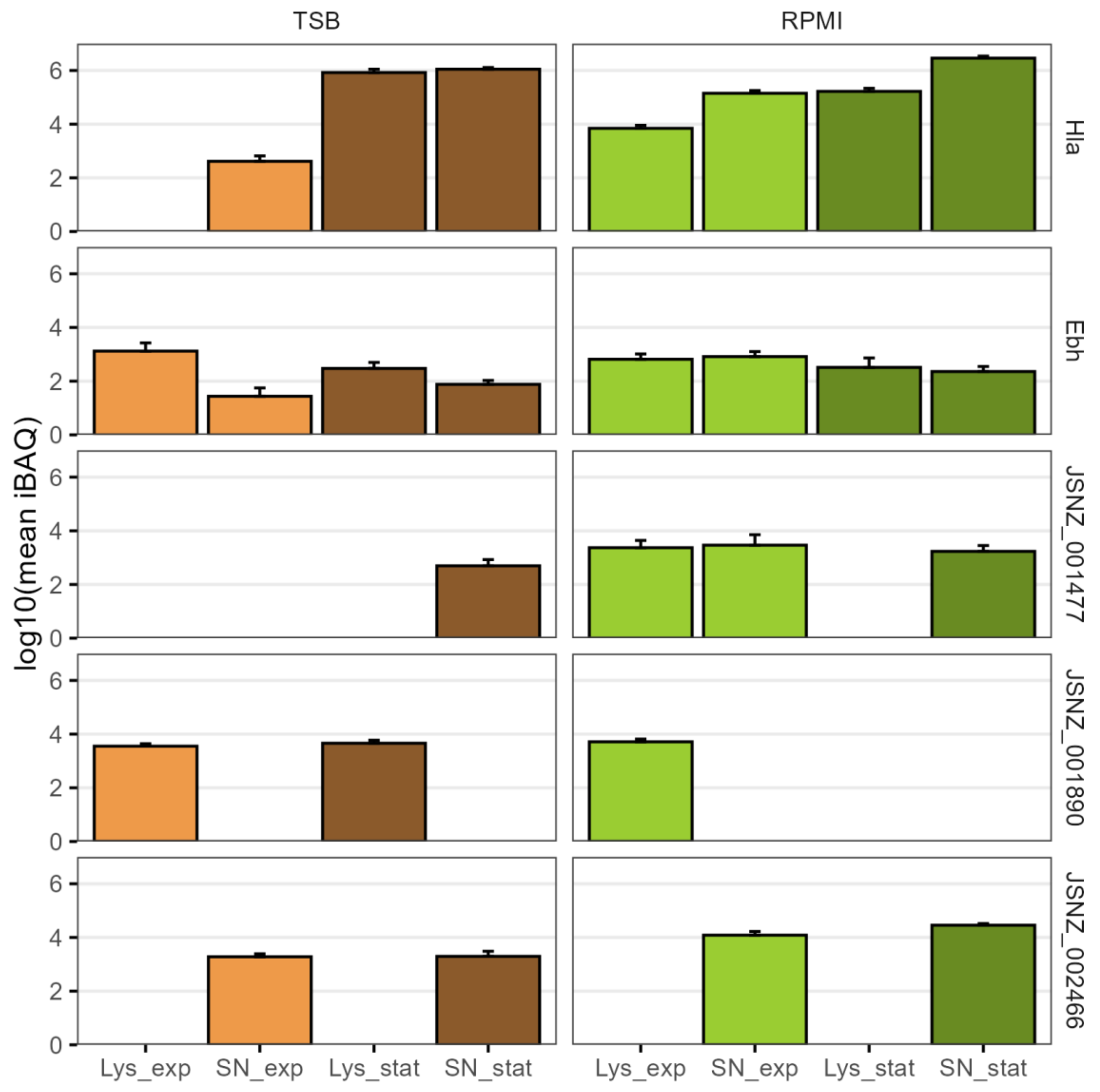


**Figure S7.** Abundance of detected proteins encoded by annotated pseudogenes in the JSNZ genome. iBAQs are calculated based on four bioreplicates, standard deviation is shown by error bars.

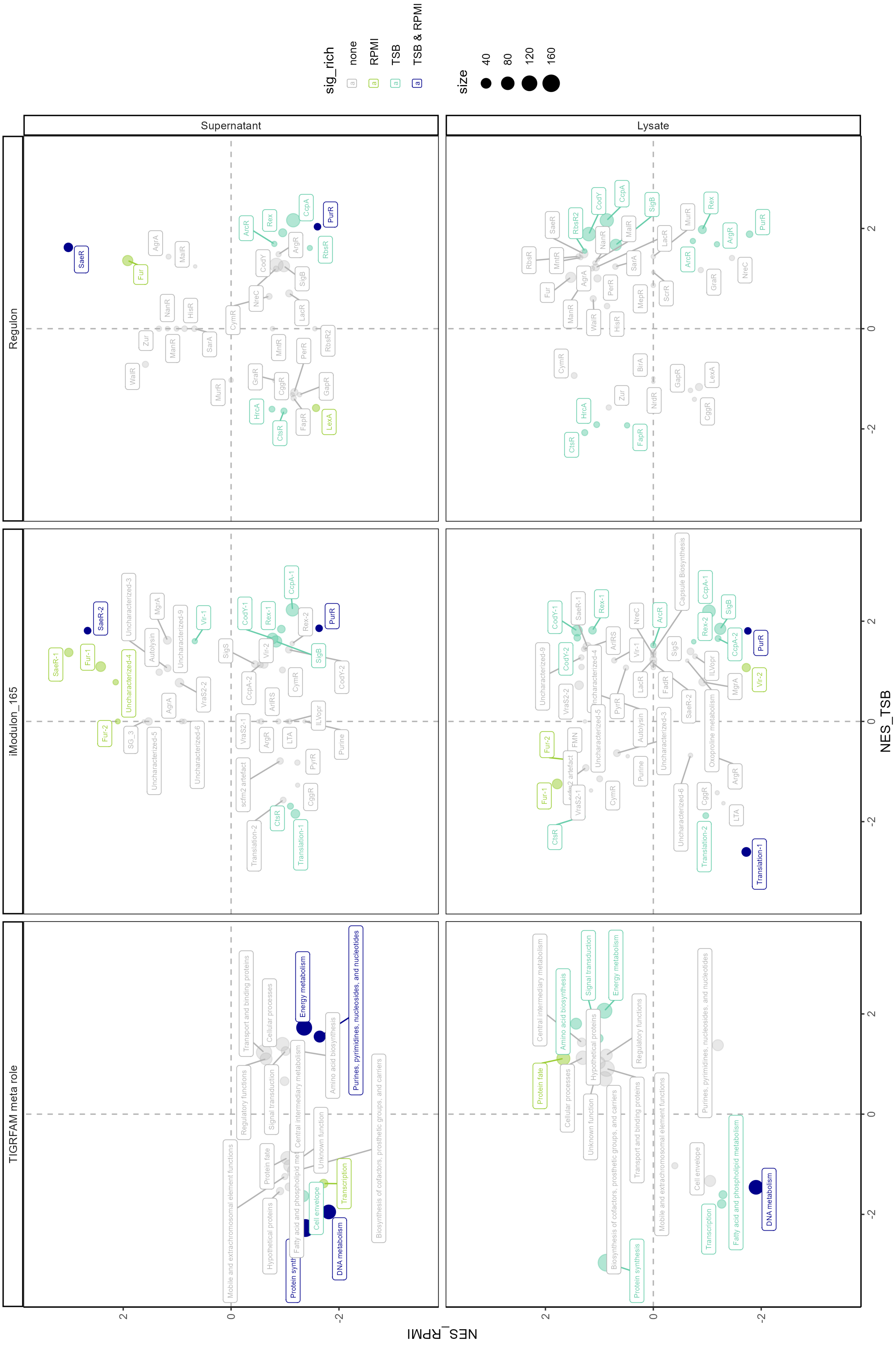

Regulon
iModulon_165
TIGRFAM meta role
Supernatant
Lysate
NES_TSB
NES_RPMI
sig_rich
none
RPMI
TSB
TSB & RPMI
size
40
80
120
160


Figure S8

**Figure S8.** Enrichment of protein sets comparing growth phases. GSEA statistics for proteome alteration from exponential to stationary growth phase per medium. Proteins were grouped to sets according to TIGRAFM meta role (left), assigned iModulon (center), and regulon (right), respectively. Points are coloured based on significant enrichment (adjusted p-value <0.05) in one or both respective comparisons, point size is relative to protein set size. Legends are centrally collected. Explanatory, schematic plots are included to aid data interpretation. This figure is the fully annotated version of Fig. 4A from the main text. NES = normalized enrichment score.

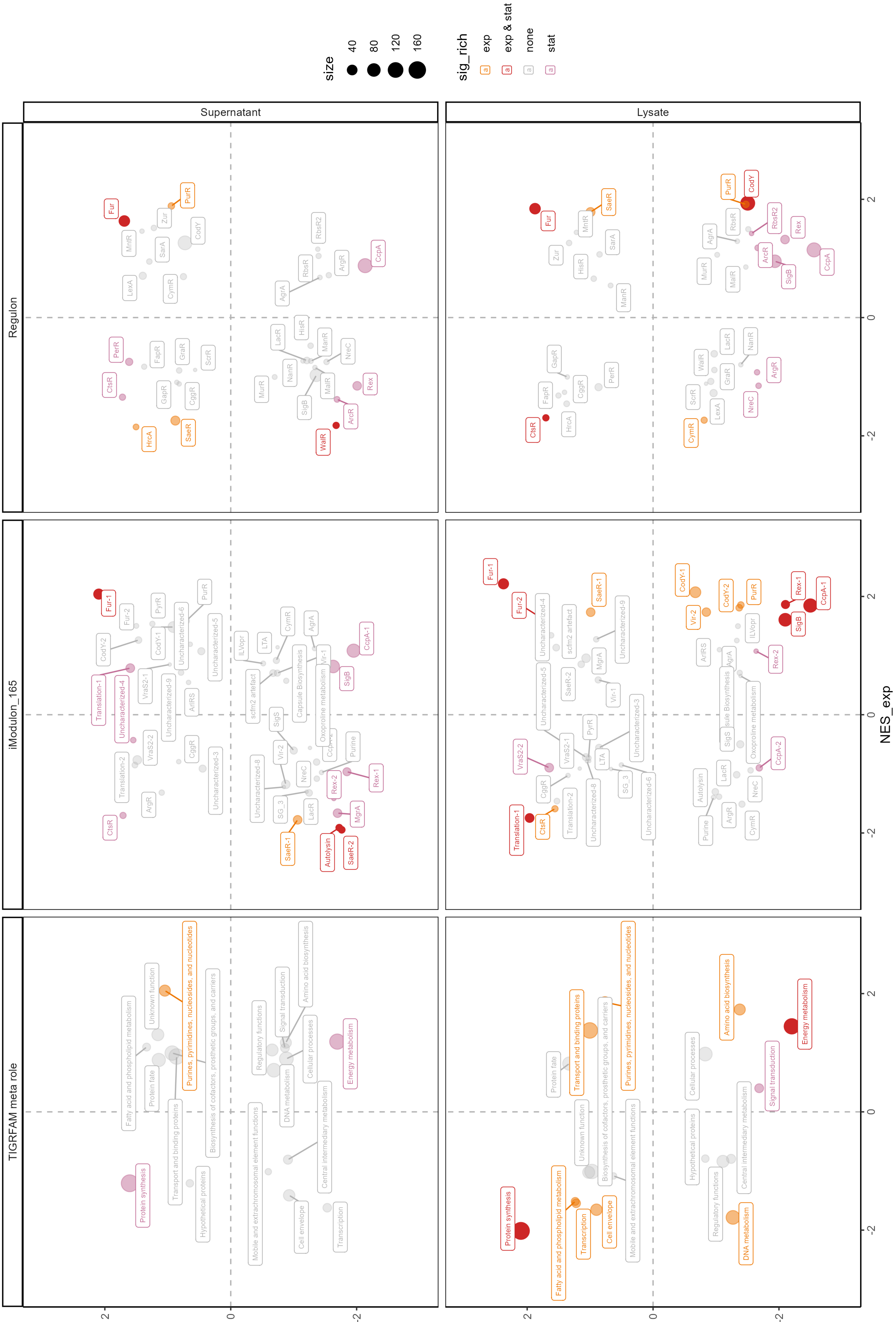

Regulon
iModulon_165
TIGRFAM meta role
Supernatant
Lysate
NES_exp
NES_stat
size
40
80
120
160
sig_rich
exp
exp & stat
none
stat


Figure S9

**Figure S9.** Enrichment of protein sets comparing media. GSEA statistics for relative proteome alteration from TSB to RPMI per growth phase. Proteins were grouped to sets according to TIGRAFM meta role (left), assigned iModulon (center), and regulon (right), respectively. Points are coloured based on significant enrichment (adjusted p-value <0.05) in one or both respective comparisons, point size is relative to protein set size. Legends are centrally collected. Explanatory, schematic plots are included to aid data interpretation. This figure is the fully annotated version of Fig. 4B from the main text. NES = normalized enrichment score.

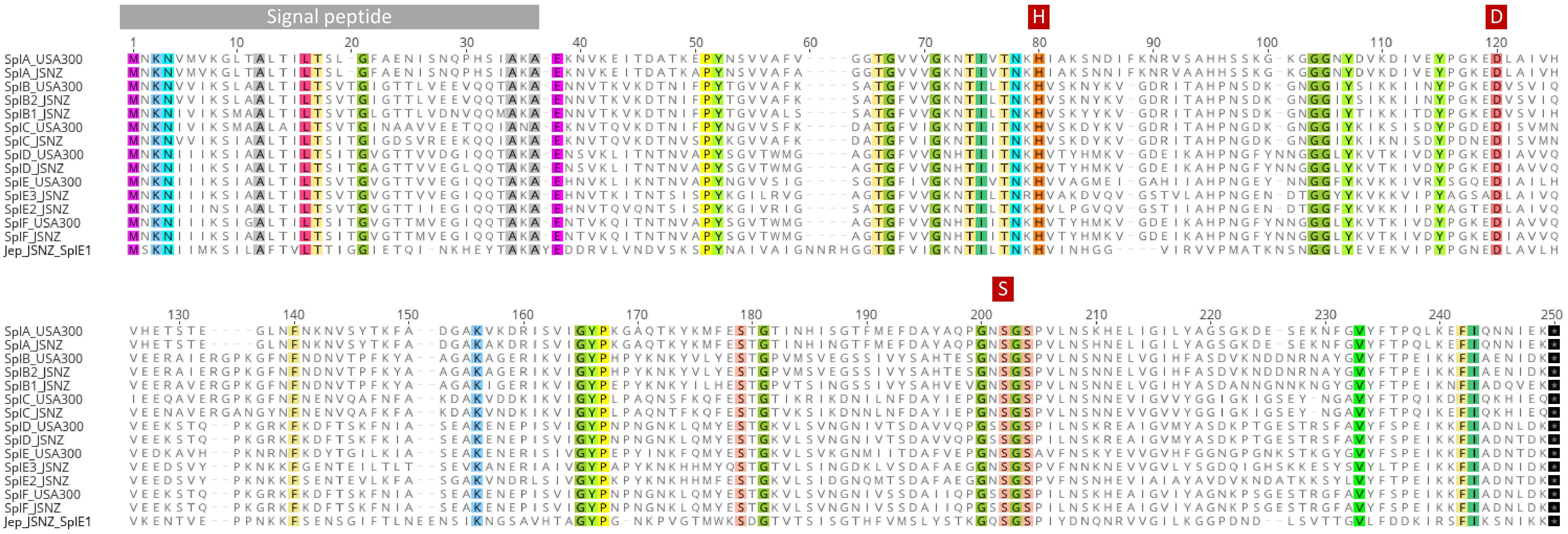

**Figure S10.** Protein sequence alignment of Jep (bottom sequence) to the Spls encoded in JSNZ and USA300_FPR3757. Conserved amino acids are coloured, signal peptide (grey) and catalytic triade (red) are indicated above the sequences. The alignment is directly visualized in Geneious Prime.

**Supplementary Tables**

Table S1. Summary of sequencing read statistics.

| | Raw reads | Trimmed reads |
|---|---|---|
| ONT | | |
| n | 102868 | 62643 |
| Total length | 726.4 Mb | 653.8 Mb |
| Mean length | 7.1 kb | 10.4 kb |
| Median length | 4.0 kb | 7.2 kb |
| N 50 | 12.6 kb | 14.2 kb |
| Min length | 43 bp | 2,468 bp |
| Max length | 139.0 kb | 139.0 kb |
| coverage | | 240.4x |
| Illumina (paired reads) | | |
| n | 5641601 | 5579151 |
| length | 2x 150 bp | 2x 36 bp to 150 bp |
| coverage | | ca. 600x |

Table S2. Reference integrase sequences used for phage classification according to Goerke *et al.* (2009).

| Class | Exemplary phage | NCBI accession |
|---|---|---|
| Sa1int | *Staphylococcus* phage phiETA | NP_510895.1 |
| Sa1int | *Staphylococcus* phage 55 | YP_240491.1 |
| Sa2int | *Staphylococcus* phage PVL | NP_058467.1 |
| Sa3int | *Staphylococcus* phage phi 13 | NP_803356.1 |
| Sa4int | *Staphylococcus* phage phiSauS-IPLA35 | YP_002332364.1 |
| Sa5int | *Staphylococcus* phage 29 | YP_240566.1 |
| Sa6int | *Staphylococcus* phage 88 | YP_240703.1 |
| Sa7int | *Staphylococcus* phage 52A | AAX91804.1 |
| Sa8int | *Staphylococcus* phage 53 | YP_239679.1 |
| Sa9int | *Staphylococcus* phage vB_SauS-phiIPLA88 | YP_002332477.1 |
| Sa10int | *Staphylococcus* phage 96 | AAX91428.1 |
| Sa11int | *Staphylococcus* phage 37 | AAX91273.1 |
| Sa12int | *Staphylococcus* phage EW | YP_240184.1 |

Table S3. Information on reversed phase liquid chromatography (RPLC).

| | |
|---|---|
| Instrument | Ultimate 3000 RSLC (Thermo Scientific) |
| Trap column | 75 µm inner diameter, packed with 3 µm C18 particles (Acclaim PepMap100, Thermo Scientific) |
| Analytical column | Accucore 150-C18, (Thermo Fisher Scientific)<br>25 cm x 75 µm, 2,6 µm C18 particles, 150 Å pore size |
| Buffer system | binary buffer system consisting of 0.1% acetic acid in HPLC-grade water (solvent A) and 100% ACN in 0.1% acetic acid (solvent B) |
| Flow rate | 300 nl/min |
| Gradient | 0min-2% B<br>2min-5% B<br>10min-7% B<br>70min-25% B<br>75min-40% B<br>77min-90% B<br>83min-90% B<br>85min-2% B<br>95min-2% B |
| Column oven temperature | 40°C |

Table S4. Information on data independent analysis (DIA) mass spectrometry.

| | |
|---|---|
| Instrument | Orbitrap Exploris$^{TM}$ 480 |
| Electrospray | Nanospray Flex$^{TM}$ Ion Source |
| Operation mode | data-independent |
| **Full Scan Properties** | |
| MS scan resolution | 120,000 |
| AGC target | 3e6 (300%) |
| Maximum ion injection time for the MS scan | 60 ms |
| Scan range | 350 to 1200 m/z |
| Microscans | 1 |
| Polarity | positive |
| RF Lens | 50% |
| Spectra data type | profile |
| **DIA Properties (MS2)** | |
| Resolution | 30,000 |
| Maximum ion injection time for the MS/MS scans | auto |
| Normalised AGC target | 3e6 (3000%) |
| Spectra data type | profile |
| Microscans | 1 |
| Isolation window | 66 |
| Isolation window width | 13 m/z |
| Window overlay | 2 m/z |
| Fixed first mass | 200 |
| HCD collision energy | 30%, normalised |
